# Lipid bilayer thinning near a ubiquitin ligase selects ER membrane proteins for degradation

**DOI:** 10.1101/2025.10.31.685944

**Authors:** Rudolf Pisa, Tom A. Rapoport

**Affiliations:** Department of Cell Biology, Harvard Medical School, and Howard Hughes Medical Institute; 240 Longwood Avenue, Boston, MA 02115, USA

## Abstract

Misfolded or unassembled membrane proteins in the endoplasmic reticulum (ER) are polyubiquitinated, translocated into the cytosol, and degraded by the proteasome, a poorly understood process that is conserved in all eukaryotes. Here, we use *S. cerevisiae* to elucidate how ER membrane proteins are selected for degradation. We show that hydrophilic residues in a trans-membrane (TM) segment cause the TM to partition into a thinned membrane region next to the ubiquitin ligase Hrd1, which then leads to substrate polyubiquitination and degradation. In the case of single-pass membrane proteins, the Hrd1-associated Der1 protein contributes to partitioning and degradation. In contrast, multi-pass proteins require Hrd1 to function on its own. Our results provide a general mechanism by which ER membrane proteins are targeted for degradation.

---

Many proteins are initially translocated into the ER lumen or integrated into the ER membrane by the Sec61 translocon and associated proteins ^1,2^. They subsequently undergo quality control, such that only correctly folded proteins are moved on along the secretory pathway or become resident in the ER. If a protein does not reach its native folded state or lacks a partner with which it normally forms a complex, it is ultimately degraded by a process called ER-associated protein degradation (ERAD) ^3^. During ERAD, luminal or membrane proteins are polyubiquitinated, moved into the cytosol, and finally degraded by the proteasome (for reviews, see ^4–9^). ERAD is conserved in all eukaryotic cells and alleviates cytotoxic stress imposed by protein misfolding. It is implicated in numerous diseases ^5,10^.

The mechanism by which misfolded luminal proteins are degraded (ERAD-L) is reasonably well understood (for review, see ^4^). In *S. cerevisiae*, ERAD-L is mediated by the Hrd1 complex, which contains the RING-finger ubiquitin ligase Hrd1 and four associated proteins (Hrd3, Usa1, Der1, Yos9). Yos9 and Hrd3 jointly create a luminal binding site that recognizes misfolded substrates. Subsequent substrate translocation into the cytosol is mediated by Hrd1 and the associated rhomboid-like protein Der1. The substrate inserts from the luminal side as a loop, such that one part of the hairpin moves through a lateral gate in Hrd1, and the other through a lateral gate in Der1 ^11,12^. The tip of the loop is postulated to move through a thinned membrane region located between the two lateral gates. The concept of local membrane thinning is based on a cytosolic cavity in the Hrd1 structure, the indentation of the detergent micelle in cryo-electron microscopy (EM) density maps, and molecular dynamics simulations ^11,13^. Membrane thinning likely reduces the energy barrier for translocation of a polypeptide through the phospholipid bilayer, a new paradigm that might apply to other systems, such as protein import into mitochondria ^14^. Once a suitable lysine residue has emerged on the cytosolic side, a polyubiquitin chain is attached by Hrd1, and the modified protein is pulled out of the membrane by the Cdc48 ATPase and its cofactors. Finally, the protein is transferred to the proteasome for degradation. All components have homologs in every eukaryotic organism, suggesting that the mechanism of ERAD-L is highly conserved.

ER membrane protein degradation is by contrast poorly understood. Specifically, it is unclear how membrane proteins are recognized as being misfolded or unassembled, and how they are targeted for degradation. Most studies have used either substrates that are multi-pass membrane proteins, whose folding status is difficult to assess, or membrane proteins that have an additional misfolded luminal or cytosolic domain, which precludes easy interpretation ^8,9^. Substrate selection is likely mediated by ubiquitin ligases, because polyubiquitination is the signal for degradation. In *S. cerevisiae*, all ER luminal proteins use the Hrd1 ligase, whereas membrane proteins employ three different ligases: Hrd1, Doa10, or Asi1-3 ^8,9,15^. Each of these enzymes has a distinct substrate specificity, although they can sometimes overlap. Hrd1 seems to be the most commonly used ligase. All three ligases might additionally facilitate polypeptide movement through the membrane. This function may also be performed by the Der1 homolog Dfm1 ^16^, but this protein likely acts downstream of a ubiquitin ligase that recognizes the substrate. As in ERAD-L, all polyubiquitinated membrane proteins are eventually extracted from the membrane by the Cdc48 ATPase complex and passed on to the proteasome ^15^.

Here we have addressed how ER membrane proteins are recognized and targeted for degradation by the Hrd1 complex. Starting with single-pass membrane proteins, we systematically varied the amino acid sequence of the TM segment. The results show that TMs with charged or hydrophilic amino acids in the middle of the membrane or the cytosolic leaflet of the lipid bilayer cause Hrd1-dependent degradation. These TMs partition into the locally thinned membrane region adjacent to the lateral gate of Hrd1, which leads to polyubiquitination in the cytosolic tail and subsequent substrate degradation. Der1 also contributes to membrane thinning, as demonstrated for single-pass model proteins and an endogenous substrate that normally forms a complex with a partner protein. In contrast, misfolded multi-pass membrane proteins do not depend on Der1, as they can only come close to Hrd1 when Der1 has moved away. Crosslinking data and a cryo-EM structure show that the dissociation of Der1 allows a second Hrd1 molecule to become a degradation target. Together, our results provide a general mechanism by which misfolded or unassembled ER membrane proteins are targeted for degradation.

## Results

### Degradation of single-pass proteins by hydrophilic residues in the TM

We first focused on single-pass membrane proteins, as they allow us to identify features of the TM segment that trigger substrate degradation. The model substrates are type I membrane proteins (**Fig. 1a**). They contain a cleavable signal sequence (ss) that places the well-folded mNeonGreen (mNG) domain and a hemagglutinin (HA) tag into the ER lumen. Following the TM segment, also called “stop-transfer signal”, is a cytoplasmic tail that ends with an ER retention signal (KKXX). This signal ensures that any protein degradation occurs by ERAD. The model proteins were expressed from a plasmid together with cytosolic mCherry (mCh), with a ribosomal skipping sequence in between. mCh serves as an internal control for the expression of the constructs.

**Fig. 1.**
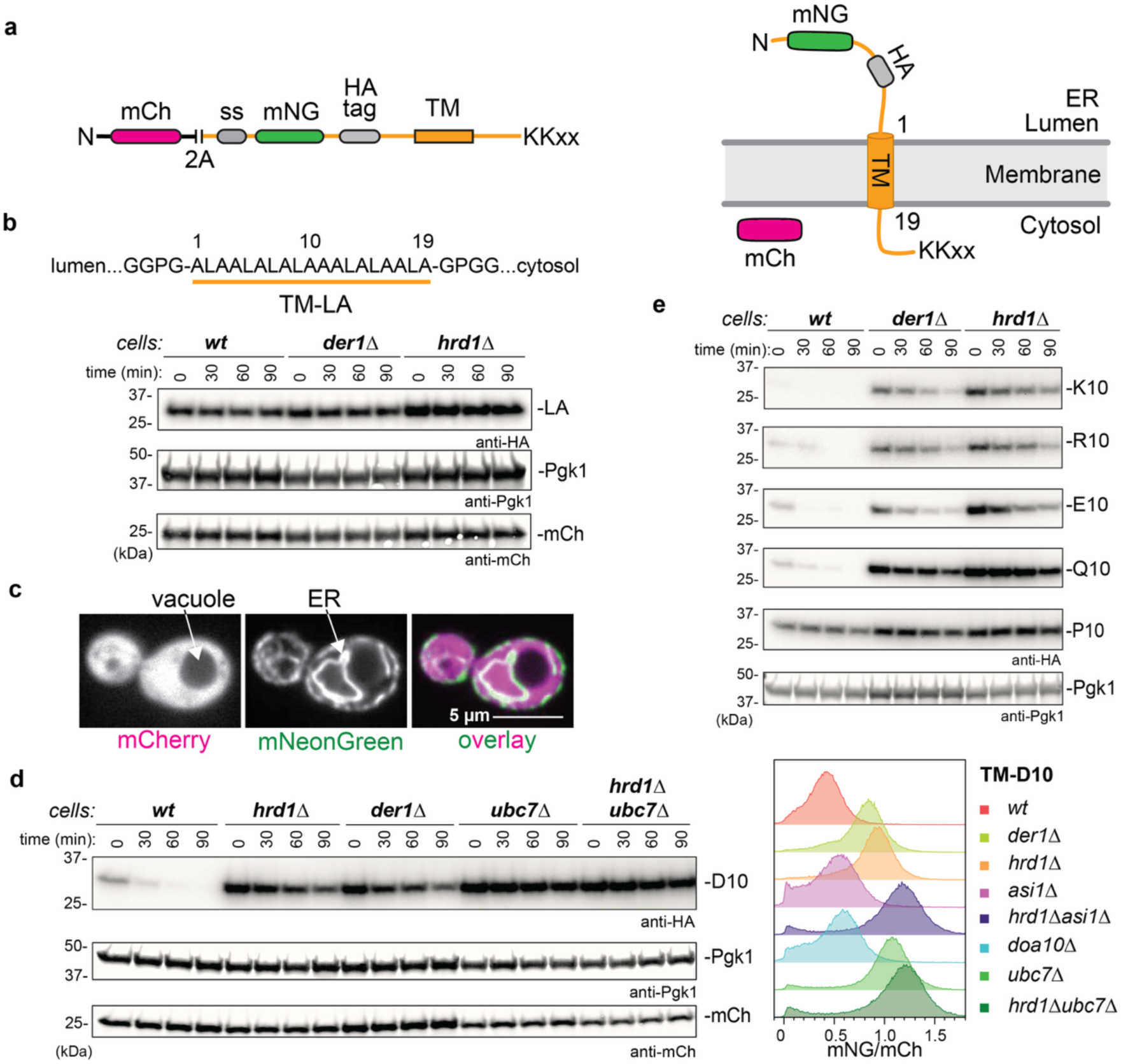
ERAD induced by hydrophilic amino acids in the middle of a TM. (a) Reporter constructs coding for cytosolic mCherry (mCh), a 2A ribosome skip site, and a type-I ER membrane protein were expressed in *S. cerevisiae*. The membrane protein contains a signal sequence (ss) that places mNeonGreen (mNG) and a HA tag into the ER lumen, followed by a trans-membrane (TM) segment and a cytosolic tail containing an ER retention signal (KKXX) (scheme on the right). (b) An idealized hydrophobic TM segment (LA) was placed into the reporter construct shown in (a). Cycloheximide-chase experiments were performed with wild-type cells (wt), or cells lacking Der1 or Hrd1, and the samples were analyzed by SDS-PAGE and immunoblotting with anti-HA antibodies. Immunoblotting for phosphoglyceratekinase (Pgk1) and mCh served as loading controls. (c) Yeast cells expressing the LA construct were analyzed by fluorescence microscopy. (d) Left: As in (b), but with a TM in which Ala at position 10 was replaced by Asp. The degradation of this D10 construct was analyzed in the indicated yeast strains. Right: Flow cytometry analysis of the indicated cells transformed with the D10 construct. (e) As in (b), but with the indicated amino acids at position 10. Loading control for P10 construct is shown.

It is difficult to identify degradation-causing amino acids in TMs of naturally occurring single-pass proteins, because their TMs are diverse in sequence and their boundaries are often ill defined. We therefore used idealized TMs that are based on a construct of 19 amino acids, composed of 7 Leu and 12 Ala residues, flanked by GPGG…GPGG linkers (LA construct; **Fig. 1b**). Such idealized TMs have been extensively studied in the context of protein translocation ^17–20^. The 7 Leu residues ensure that the TM is sufficiently hydrophobic to allow it to function as a stop-transfer sequence and to cause its stable integration into the phospholipid bilayer ^17,18^. Consistent with previous reports ^18^, the LA construct sedimented exclusively with membranes in cell fractionation experiments (**Extended Data Fig. 1a**). Importantly, the hydrophobic TM in the LA construct resulted in a completely stable protein, as shown by cycloheximide-chase experiments (**Fig. 1b**). Furthermore, fluorescence microscopy indicated that the protein localizes to the ER, whereas co-expressed mCherry was found in the cytosol (**Fig. 1c**).

Next, we replaced the Ala residue with Asp at position 10 of the TM (D10 construct). The protein was readily degraded in wild-type yeast cells, but considerably stabilized in strains lacking Hrd1 or Der1 (**Fig. 1d**). Complete stabilization was observed in a strain lacking the ubiquitin-conjugating E2 enzyme Ubc7 or both Hrd1 and Ubc7 (**Fig. 1d**), consistent with Ubc7 being the major E2 enzyme for all ER-localized ubiquitin ligases ^21^. Measurement of the steady-state levels by flow cytometry confirmed that the degradation of the D10 construct depends on Hrd1 and Der1 (**Fig. 1d**; right panel). In contrast, the absence of Asi1-3, components of the Asi-ligase complex ^21^, the ubiquitin ligase Doa10 ^22^, the autophagy component Atg8, or the nuclear ubiquitin ligase San1 had only small effects (**Extended Data Fig. 1b**). The D10 construct was degraded in a Cdc48-dependent manner: it was unstable in a *cdc48-3* mutant strain at the permissive, but not non-permissive, temperature (**Extended Data Fig. 1c**). As expected from the calculated negative ΔG value of membrane integration ^17^, the D10 substrate exclusively fractionated with membranes when its degradation was prevented by *hrd1* deletion (**Extended Data Fig. 1a**). These results show that an Asp residue in the middle of the TM causes Hrd1- and Der1-dependent ERAD of the model protein.

Several other hydrophilic amino acid residues (Lys, Arg, Glu, Gln) at position 10 of the TM also caused degradation of the model protein (**Fig. 1e; Extended Data Fig. 1d**). Introducing a Pro residue at position 10 left the model protein stable, suggesting that perturbation of the TM helix is not a signal for degradation. Again, all proteins were stabilized in the absence of Der1 or Hrd1, although Hrd1 always had a stronger effect. By contrast, the absence of Pep4, a key vacuolar protease, did not slow degradation (**Extended Data Fig. 1d**). Deleting *asi1* also had little effect, but deleting *asi1* and *hrd1* together generally resulted in more stabilization than *hrd1* deletion alone (**Extended Data Fig. 1d**), suggesting that the Asi1 complex makes a small contribution to the degradation of the model substrates.

## Hydrophilic residues placed in the cytosolic leaflet are destabilizing

Next, we placed a hydrophilic residue at different positions of the TM (**Fig. 2a**). An Asp residue introduced close to the luminal side of the TM (D1 or D4) caused some degradation, but much less than an Asp residue near the cytosolic side (D10, D16, D19) (**Fig. 2a**). As shown for D19, the degradation was again dependent on Der1 (**Extended Data Fig. 2a**).

**Fig. 2.**
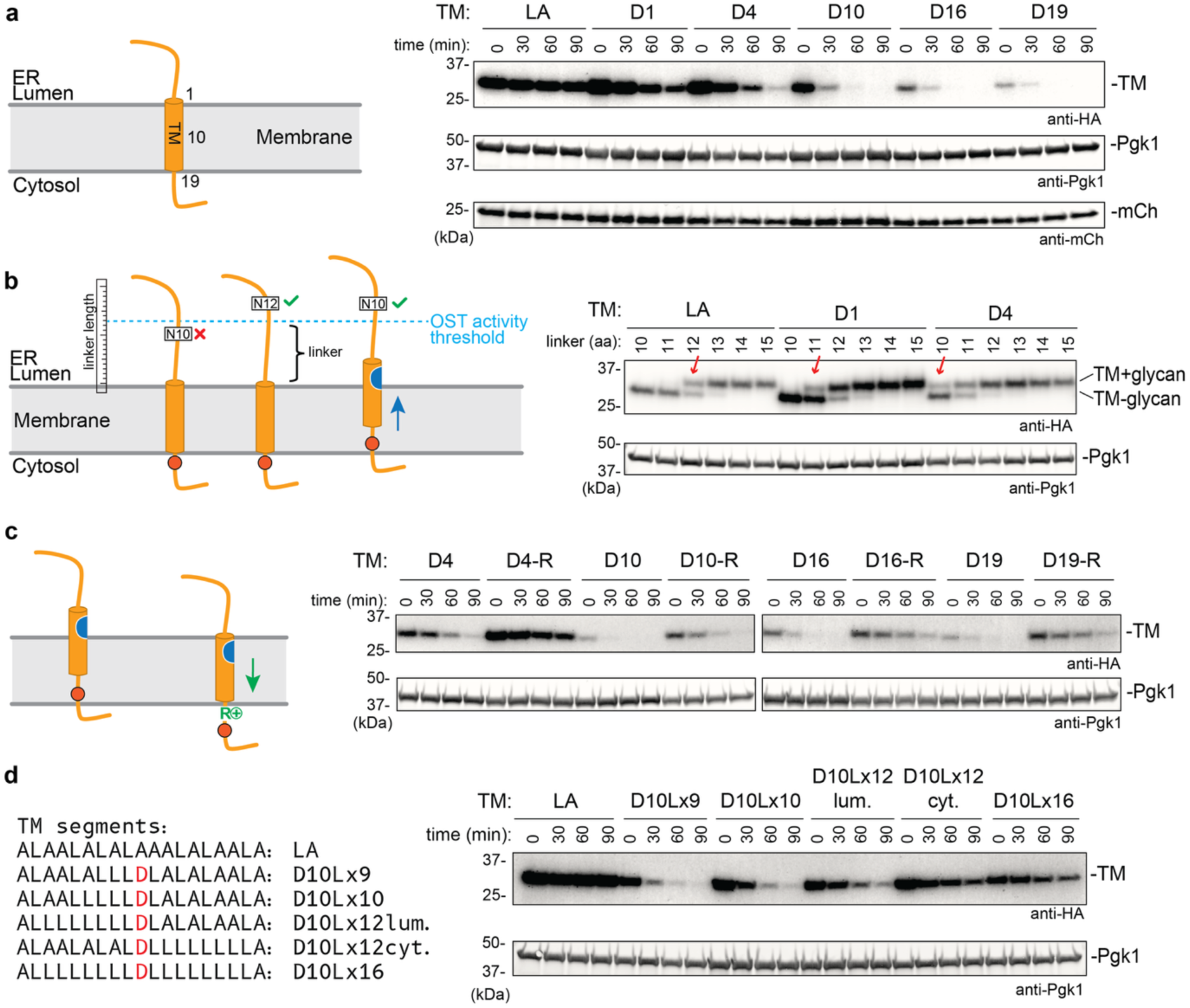
ERAD depends on the position of a hydrophilic amino acids in the TM. (a) HA-tagged constructs containing the hydrophobic LA segment or TMs with Asp residues at the indicated positions were tested for their degradation by cycloheximide-chase experiments. The samples were subjected to SDS-PAGE and immunoblotting with anti-HA antibodies. Immunoblotting for phosphoglyceratekinase (Pgk1) and mCh served as loading controls. (b) Scheme illustrating that a TM with a hydrophilic residue (in blue) can slide into the lumen and pull instead residues from the cytosol (red dot) into the membrane. The minimal distance from the lumenal side required for N-glycosylation by oligosaccharyl transferase (OST) is 12 residues (red cross: no modification; green check sign, modification). HA-tagged LA, D1, or D4 constructs with the indicated N-glycosylation sites at different luminal positions were expressed in cells lacking Hrd1 and Asi1. The attachment of a glycan was determined by the mobility shift in immunoblots (right panel). The red arrows point to minimal linker length at which glycosylation was observed. (c) As in (a), but where indicated, the constructs contained the RPRR sequence instead of the GPGG linker at the cytosolic end of the TM to prevent sliding of the TM into the ER lumen. (d) As in (a), but with Asp at position 10 (in red) and replacement of some Ala residues with Leu. Note that increasing the hydrophobicity in the cytosolic leaflet slows degradation.

Because some degradation was observed even with the D1 and D4 constructs, we wondered if these residues could slide into the ER lumen and pull instead residues of the cytosolic tail into the membrane (**Fig. 2b**, scheme). Sliding was indeed confirmed by placing an N-glycosylation site at different positions of the luminal region of the constructs (**Fig. 2b**). With the hydrophobic LA construct, modification was observed when the glycosylation site was at least 12 residues away from the luminal end of the TM, consistent with the known minimum distance for modification by the oligosaccharyl transferase ^23^. With the D1 and D4 constructs, however, glycosylation was observed at shorter distances (11 and 10 residues, respectively) (**Fig. 2b**). To prevent sliding, we replaced the cytosolic GPGG linker with the RPRR sequence in the D4 construct (see scheme in **Fig. 2c**). Pulling the Arg residues of this sequence into the membrane would be energetically unfavorable. Indeed, the minimal glycosylation distance was increased (**Extended Data Fig. 2b**) and the D4 residue was no longer destabilizing, whereas the D10 and D19 residues continued to be degradation signals (**Fig. 2c).** Control experiments showed that an RPRR linker introduced at the equivalent position of the LA construct had no effect (**Extended Data Fig. 2c**). Taken together, these results suggest that TMs exposing hydrophilic residues to phospholipids in the cytosolic leaflet cause ERAD.

The model is supported by experiments in which we added Leu residues on either the luminal or cytosolic side of the TM in the D10 construct (**Fig. 2d**). Making the luminal side more hydrophobic had no effect, but introducing a stretch of consecutive Leu residues on the cytosolic side inhibited degradation, consistent with these residues facilitating the placement of D10 into lipids. Thus, hydrophilic residues inside the membrane can be tolerated in stop-transfer signals because the overall hydrophobicity of the TM still allows membrane integration, but they target the protein for degradation.

The identified degradation rules also apply to naturally occurring TMs. A TM from the multi-pass protein Pdr5 that contains a central Asn residue ^24^ triggered rapid Hrd1- and Der1-dependent degradation when placed into our reporter construct (**Extended Data Fig. 2d**). When the Asn residue was mutated to Leu, the protein remained stable (**Extended Data Fig. 2d**). Likewise, the completely hydrophobic TM from the Wsc1 protein did not cause degradation (**Extended Data Fig. 2e**).

We were surprised that the absence of Der1 retarded the degradation of our model substrates, because Der1 had no effect in previous experiments with multi-pass proteins ^15^. One possible explanation is that single-pass proteins move all the way into the ER lumen and become Der1-dependent ERAD-L substrates. However, this possibility was excluded by introducing an N-glycosylation site into the cytosolic tail of our model substrates (**Extended Data Fig. 2f**). In the case of D4 and D10, a small population was glycosylated, but the modified species was relatively stable during the cycloheximide chase. For D16 and D19, no glycosylated species were observed. Thus, these proteins do not seem to move into the ER lumen before being degraded. Placing the RPRR linker at the cytosolic end of the TM to prevent its sliding into the lumen also did not abolish Der1-dependency of degradation (**Fig. 2c and Extended Data Fig. 2g**). We therefore conclude that single-pass membrane proteins are substrates that use both Hrd1 and Der1 for their degradation.

## Degradation-causing residues are energetically unfavorable in the bilayer

Next, we performed a more systematic and quantitative analysis to determine which amino acids are destabilizing. We designed a library in which all possible amino acids were randomly incorporated at position 10 of the TM (**Fig. 3a**). To selectively turn off the synthesis of the model proteins, while leaving all other proteins unaffected, we introduced three tetracycline aptamers at the 5’end of the coding sequence ^25^. Addition of tetracycline indeed caused Hrd1-dependent degradation of the D10 construct, similarly to the addition of cycloheximide (**Extended Data Fig. 3a**). Cells expressing TM constructs with randomized position 10 were subjected to cell sorting on the basis of the ratio of mNG over mCh (**Fig. 3a, b**). The populations with the lowest and highest ratios (**Fig. 3b**) were subjected to next-generation DNA sequencing and the encoded amino acids were ranked according to their enrichment or depletion relative to the unsorted library (**Fig. 3b**). The results show that Lys, Glu, Arg, Gln, Asp, His, and Asn were depleted in the high mNG/mCh population and enriched in the low mNG/mCh population (**Fig. 3c**). Thus, all hydrophilic residues, with the exception of Ser and Thr, cause degradation. By contrast, all hydrophobic amino acids were stabilizing. The relative abundance in the high mNG/mCh population showed an excellent correlation with the free energy ΔG required to move an amino acid in a translocating polypeptide from the Sec61 translocon into the lipid phase (biological hydrophobicity scale ^17^) (**Fig. 3d**). A reasonable correlation with this scale was also seen when the depletion in the high mNG/mCh population and the enrichment in the low mNG/mCh population were both taken into account (as the ratio of the relative abundances, here called TM stability score; TMSS) (**Fig. 3e**). Because the biological hydrophobicity scale is related to the propensity of amino acids to partition from water into octanol ^17,26^, the TMSS also correlated with the ΔG associated with this transfer (**Extended Data Fig. 3b**). Thus, the destabilizing effect of an amino acid in the middle of the TM depends on how energetically unfavorable it is in the phospholipid bilayer.

**Fig. 3.**
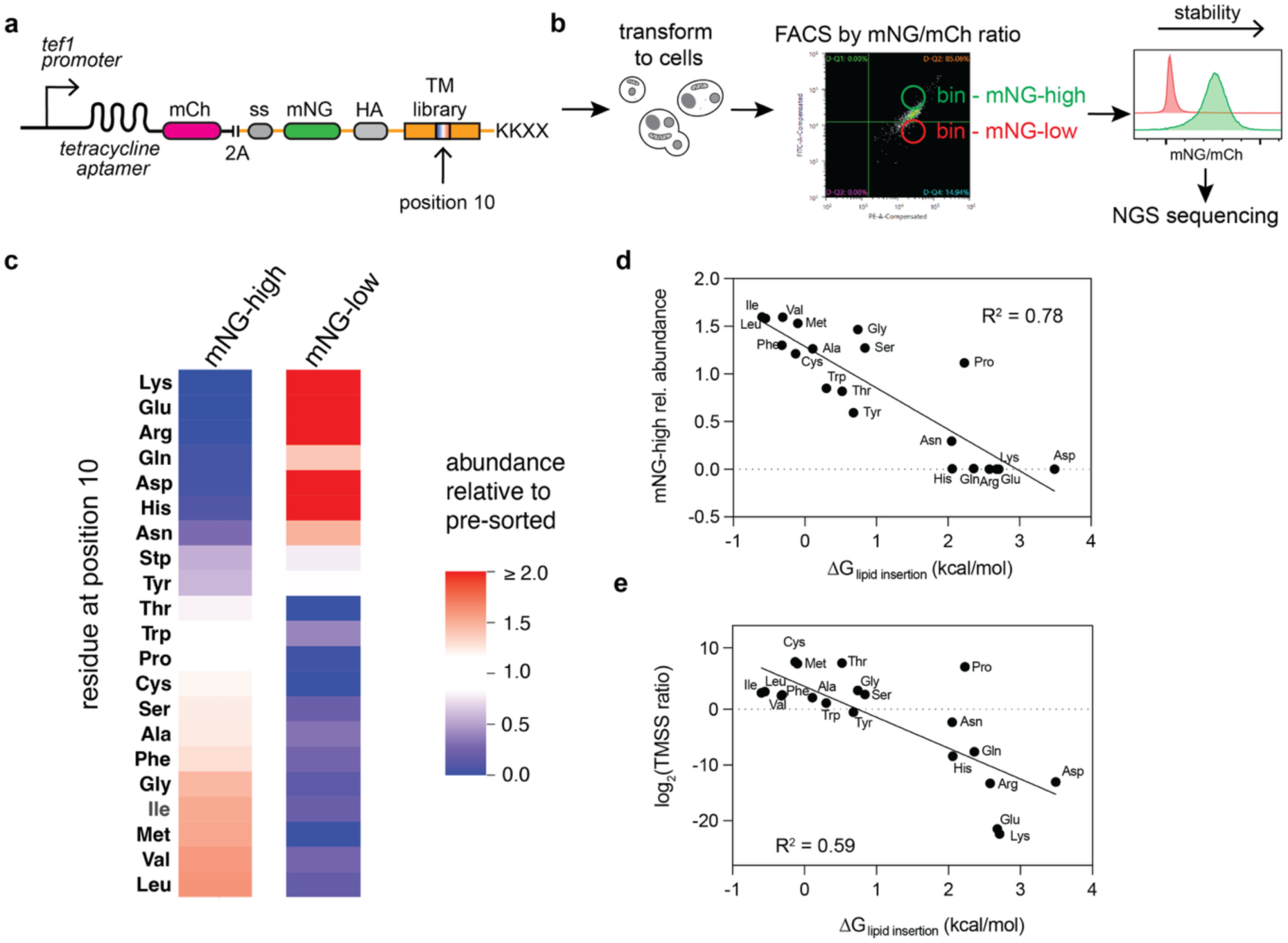
The energetic penalty of amino acid transfer into lipid determines degradation. (a) The indicated reporter construct under the control of the tef1 promoter and tetracyclin aptamers was used to create a TM library with each amino acid randomly incorporated at position 10. (b) The library was expressed in yeast cells grown in tetracyclin to turn off expression, and subjected to cell sorting (FACS) based on the mNG/mCh ratio. Cells with the highest and lowest ratio values were collected and analyzed by flow cytometry. Both pools and the unsorted library were subjected to next-generation DNA sequencing (NGS). (c) Amino acids depleted (blue) or enriched (red) in the high and low mNG/mCh cell pools, relative to their abundance in the unsorted library (abundance scale on the right). (d) The depletion of an amino acid in the high mNG/mCh population relative to its abundance in the unsorted library was plotted against the free energy of transfer of an amino acid from aqueous solution into the lipid bilayer (ΔG_lipid insertion_; biological hydrophobicity scale – values from ref. ^17^. A linear regression line was calculated from the points by least square fitting. R^2^ is the correlation coefficient. (e) A TM stability score (TMSS) was calculated as the ratio of the relative depletion in the high mNG/mCh population over the enrichment in the low mNG/mCh population. Plotted is the log_2_ of TMSS versus ΔG_lipid insertion_.

## Degradation of an orphaned endogenous single-pass substrate

Our model predicts that endogenous single-pass proteins with destabilizing hydrophilic residues in their TM can only avoid degradation when associated with a partner protein that shields such residues from the lipid environment. We identified the Erv25 protein as a candidate to test this hypothesis. Erv25 is a member of the p24 family that is involved in cargo transport between the ER and Golgi ^27^. It has Asn and Gln residues close to the predicted cytosolic end of the TM and contains an ER retrieval signal in its cytosolic tail (**Fig. 4a**). Erv25 normally associates with Emp24, another p24 family member ^27^, raising the possibility that the hydrophilic TM residues are buried in the complex. Indeed, when HA-tagged Erv25 was moderately overexpressed in wild-type cells from a CEN plasmid under the endogenous promoter, the excess over its partner Emp24 was rapidly degraded (**Fig. 4b**); the entire Erv25 population was degraded in cells lacking Emp24 (**Fig. 4b**). The protein was stabilized in the absence of Hrd1 or Der1 (**Fig. 4b**). The stabilized protein showed a time-dependent size increase, caused by mannosylation in the Golgi (data not shown), consistent with the protein cycling between the ER and Golgi.

**Fig. 4.**
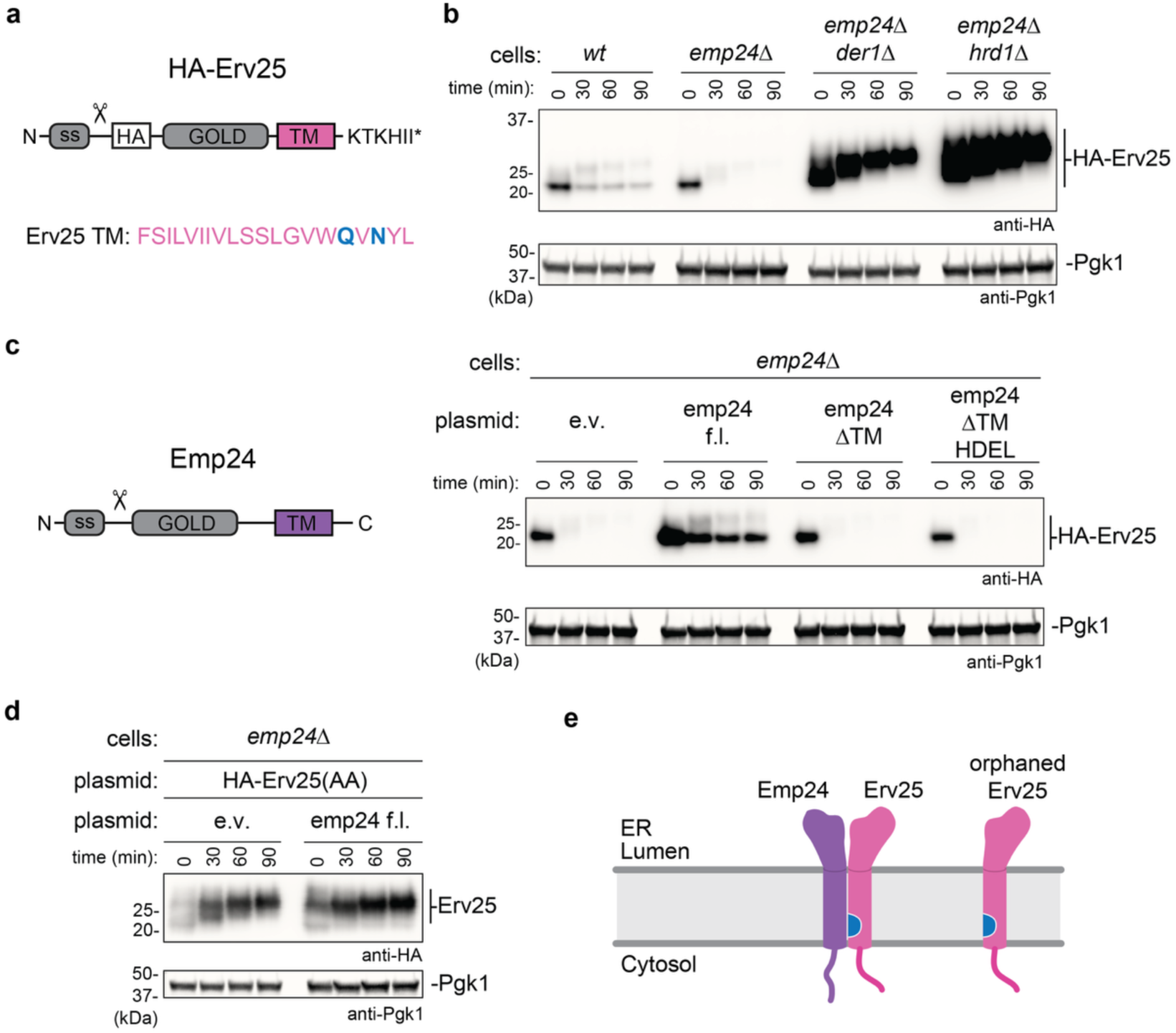
An endogenous single-pass protein undergoes ERAD when orphaned. (a) Domain structure of the HA-tagged single-pass protein Erv25. The construct contains a cleavable signal sequence (SS; cleavage site indicated by a scissor), an HA tag, a luminal GOLD domain, a TM segment, and an ER retention signal. The predicted TM has two hydrophilic residues (in blue) close to its cytosolic end. (b) HA-Erv25 was expressed in wild-type (wt) cells or cells lacking the indicated components. Cycloheximide-chase experiments were performed and the samples analyzed by SDS-PAGE and blotting with anti-HA antibodies. The time-dependent mobility shift is caused by mannosylation of HA-Erv25 in the Golgi. Immunoblotting for phosphoglyceratekinase (Pgk1) served as a loading control. (c) Left: domain structure of Emp24, the binding partner of Erv25. Right: Cycloheximide-chase experiments of HA-Erv25 expressed in cells lacking Emp24. The cells also contained either an empty vector (e.v.) or plasmids encoding full-length (f.l.) Emp24, Emp24 lacking its TM (Emp24ΔTM), or Emp24 in which the TM was replaced with an HDEL ER-retention signal (Emp24ΔTM HDEL). (d) The two hydrophilic residues in the TM of HA-Erv25 were replaced by Ala. The stability of this protein was analyzed as in (c). (e) Hydrophilic residues in the TM of Erv25 (in blue) are shielded in the Emp24-Erv25 dimer and exposed in orphaned Erv25.

HA-Erv25 was stabilized in cells containing Hrd1 and Der1 when Emp24 was co-expressed, and this stabilization could be attributed to the TM of Emp24 (**Fig. 4c**). Mutation of the two hydrophilic TM residues into Ala abolished the degradation of HA-Erv25 in the absence of Emp24 (**Fig. 4d**). Emp24 on its own has no ER retention signal and is degraded in part in vacuoles (**Extended Data Fig. 4a**). Taken together, these data show that the TM of orphaned Erv25 causes ERAD by the exposure of hydrophilic residues to the surrounding phospholipids; in the Erv25-Emp24 complex, these residues are shielded by the TM of Emp24 (**Fig. 4e**).

## Partitioning of hydrophilic TMs to the thinned membrane next to Hrd1

Given that all hydrophilic residues, including both positively and negatively charged amino acids, cause Hrd1-dependent degradation, it seemed unlikely that the ligase recognizes substrates by specific amino acid interactions. We therefore postulated that the TMs would partition into the thinned membrane region next to Hrd1, which had been visualized in previous cryo-EM structures as a cytosolic cavity next to the lateral gate of Hrd1 ^11^. TMs with charged or hydrophilic residues in the cytosolic leaflet would thus move from the energetically unfavorable lipid phase into the aqueous environment inside the cavity (**Extended Data Fig. 4b**). The increased residence time next to Hrd1 would explain why the substrates are polyubiquitinated and degraded.

We first tested whether Hrd1 indeed has a cytosolic cavity in intact ER membranes. Single Cys residues were introduced at various positions of Hrd1 and their accessibility to a membrane-impermeable modification reagent (5 kDa maleimide polyethyleneglycol, mal-PEG) was tested (**Extended Data Fig. 4c**). Positions F88C, I91C, W305C, and Q312C in the cytosolic cavity were partially modified in the absence of detergent (red arrows), and completely in the presence of detergent. Cysteines close to the ER lumen (positions N92, S98 and M297) or positions facing lipids in the cytosolic leaflet (I307 and L317) were not modified in the absence of detergent (**Extended Data Fig. 4c; 4d**). The data confirm that Hrd1 has a pronounced cytosolic cavity that extends about halfway across the membrane.

Next, we tested whether TMs with hydrophilic residues have an increased residence time next to the cytosolic cavity of Hrd1. Because such encounters are rare for membrane proteins that freely diffuse in the lipid bilayer, we used for these experiments the single-pass Hrd3 protein, which remains in proximity of Hrd1 because it is bound to Hrd1 through its luminal domain (**Fig. 5a**). If the TM has an increased residence time next to lateral gate of Hrd1, it should compete with the ERAD-L substrate CPY* and prevent its degradation. Indeed, whereas the hydrophobic TM of wild-type Hrd3 allowed efficient CPY* degradation, replacing it with a TM of Pdr5 that contains a central Asn residue caused strong inhibition (**Fig. 5b**). The hydrophobic TM of the single-pass protein Spt23 was only a weak competitor. As expected, deleting the cytosolic tail of Hrd3 (Δtail) or both the TM and the tail (ΔTM+tail) were without effect. Replacing Hrd3’s TM with the hydrophobic LA sequence also did not inhibit CPY* degradation (**Fig. 5c**), whereas TMs with Asp residues were strong competitors (**Fig. 5c**). It should be noted that all Hrd3 chimeras were expressed at levels similar to wild-type Hrd3 (**Extended Data Fig. 5**). These data suggest that hydrophilic TM segments are trapped near the lateral gate of Hrd1 for prolonged time periods thus blocking the access of the luminal substrate. Hydrophobic TMs can come close to the lateral gate but dissociate rapidly and therefore do not block the access.

**Fig. 5:**
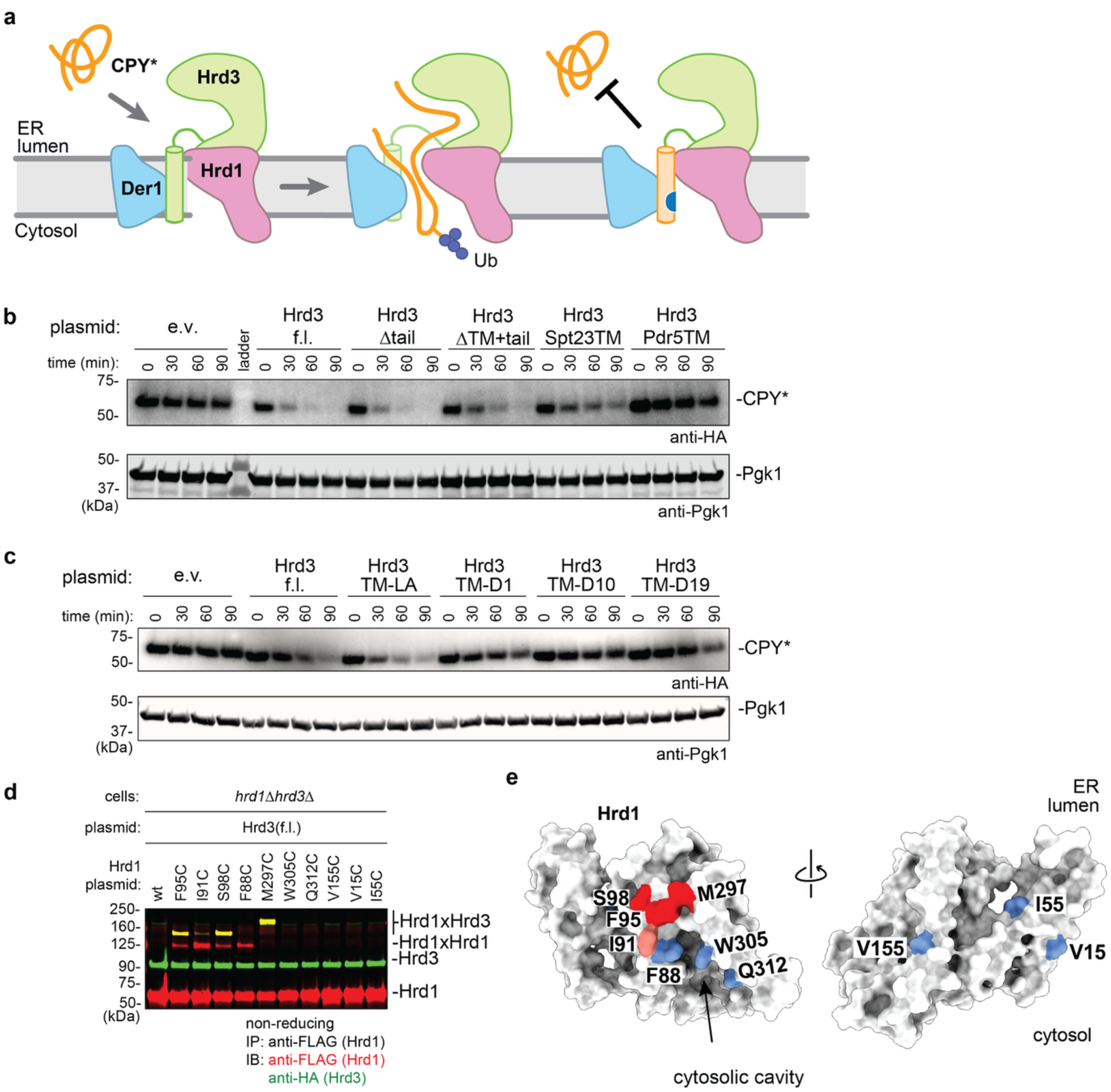
TMs with hydrophilic residues have an increased residence time near Hrd1. (a) The hydrophobic TM of Hrd3 has a short residence time near Hrd1 and therefore does not prevent loop insertion of the ERAD-L substrate CPY* into the Hrd1 complex, allowing CPY* polyubiquitination and degradation (first two panels). TMs with hydrophilic residues (blue) partition into Hrd1’s hydrophilic cavity and therefore have a long residence time, blocking CPY* loop insertion (last panel). (b) The degradation of HA-CPY* was followed by cycloheximide-chase experiments in Δhrd3 cells containing an empty vector (e.v.) or plasmids coding for full-length (f.l.) Hrd3, Hrd3 lacking its cytoplasmic tail (Δtail) or both its TM and tail (ΔTM+tail), or Hrd3 in which the endogenous TM was replaced by TMs from Spt23 (MLLFFWIPLTLVLLLCFTLS**N**LG) or Pdr5 (NYGIFICYIAF**N**YIAGVFFYWLA). Note the hydrophilic Asn residues in both TMs. Immunoblotting for phosphoglyceratekinase (Pgk1) served as a loading control. (c) As in (b), but with the endogenous TM of Hrd3 replaced with idealized TMs. (d) Disulfide crosslinking of the TM of Hrd3 with the lateral gate of Hrd1. HA-tagged Hrd3 with endogenous Cys773 in the TM (LVTMG**C**ILGIFLLSILMSTLAA) was expressed together with FLAG-tagged Hrd1 containing cysteines at the indicated positions in cells lacking Hrd3 and Hrd1. Disulfide bond formation was induced with an oxidant, and a detergent extract subjected to immunoprecipitation with FLAG antibodies. The samples were analyzed by non-reducing SDS-PAGE followed by blotting with HA- and FLAG-antibodies. (e) Space-filling model of Hrd1 with disulfide-crosslinking positions in red or pink, and non-crosslinking positions in blue.

Disulfide crosslinking experiments confirmed that the TM of Hrd3 comes close to the lateral gate of Hrd1. An endogenous Cys residue in the TM of Hrd3 (Cys 773) crosslinked efficiently to cysteines introduced at several positions at the lateral gate of Hrd1 (**Fig. 5d**; yellow bands; **Fig. 5e**). Positions outside the lateral gate did not crosslink (**Fig. 5d**; **e**). Disulfide crosslinking depended on the cysteines in both proteins (**Extended Data Fig. 5a**). The TM of Pdr5, which has an endogenous Cys at its luminal side, crosslinked to a Cys near the luminal end of Hrd1’s lateral gate (position 98), whereas the TM of Spt23, which has an endogenous Cys closer to the cytosolic end, crosslinked to a more cytosolically positioned Cys in Hrd1 (position 91) (**Extended Data Fig. 5b**). Idealized TMs containing single Cys residues at various positions also crosslinked to the lateral gate of Hrd1 (**Extended Data Fig. 5c**). Similar experiments with Der1 showed that Cys residues at positions 1-4 of an idealized TM could form disulfide crosslinks with several positions at its lateral gate (**Extended Data Fig. 5d**). Taken together, these data indicate that the TM of Hrd3 frequently encounters the thinned membrane region near Hrd1 and Der1.

Although Hrd3’s TM is in close proximity to Hrd1, Hrd3 is not degraded (**Extended Data Fig. 5a)**, because the cytosolic tail lacks a Lys residue that can be polyubiquitinated. When a Lys was introduced at two different positions of the tail, Hrd3 was undetectable and disulfide crosslinks between Hrd1 and Hrd3 disappeared (**Extended Data Fig. 5e**). These data support the idea that partitioning of a TM into the cytosolic cavity of Hrd1 and the presence of a Lys residue in the cytosolic tail are both required for substrate degradation. Consistent with this conclusion, Hrd3 in all species lacks Lys residues in the cytosolic tail (**Extended Data Fig. 5f)**, ensuring that it is not polyubiquitinated by Hrd1 despite its proximity.

## Degradation of multi-pass membrane proteins by Hrd1

Next we tested whether multi-pass membrane protein degradation is also triggered by the exposure of hydrophilic residues to lipids and their partitioning to the thinned membrane region near Hrd1. Asn residues were introduced into HA-tagged Sec61 at positions expected to face lipids inside the membrane (**Fig. 6a**), and the stability of HA-Sec61 tested with cycloheximide-chase experiments. All mutants were less stable than the wild-type protein (**Fig. 6b**) and stabilized in the absence of Hrd1 (**Extended Data Fig. 6a, b**). The degradation of the V375N mutant was not affected by the absence of Asi1 or Doa10 (**Fig. 6c**). As reported for other multi-pass membrane proteins ^15^, the absence of Der1 did not block degradation (**Fig. 6c**). Mutating residue 375 to Asp also caused Hrd1-dependent and Der1-independent degradation of HA-Sec61 (**Fig. 6d**). Thus, a single lipid-exposed hydrophilic residue in a multi-pass protein can trigger Hrd1-dependent degradation.

**Fig. 6.**
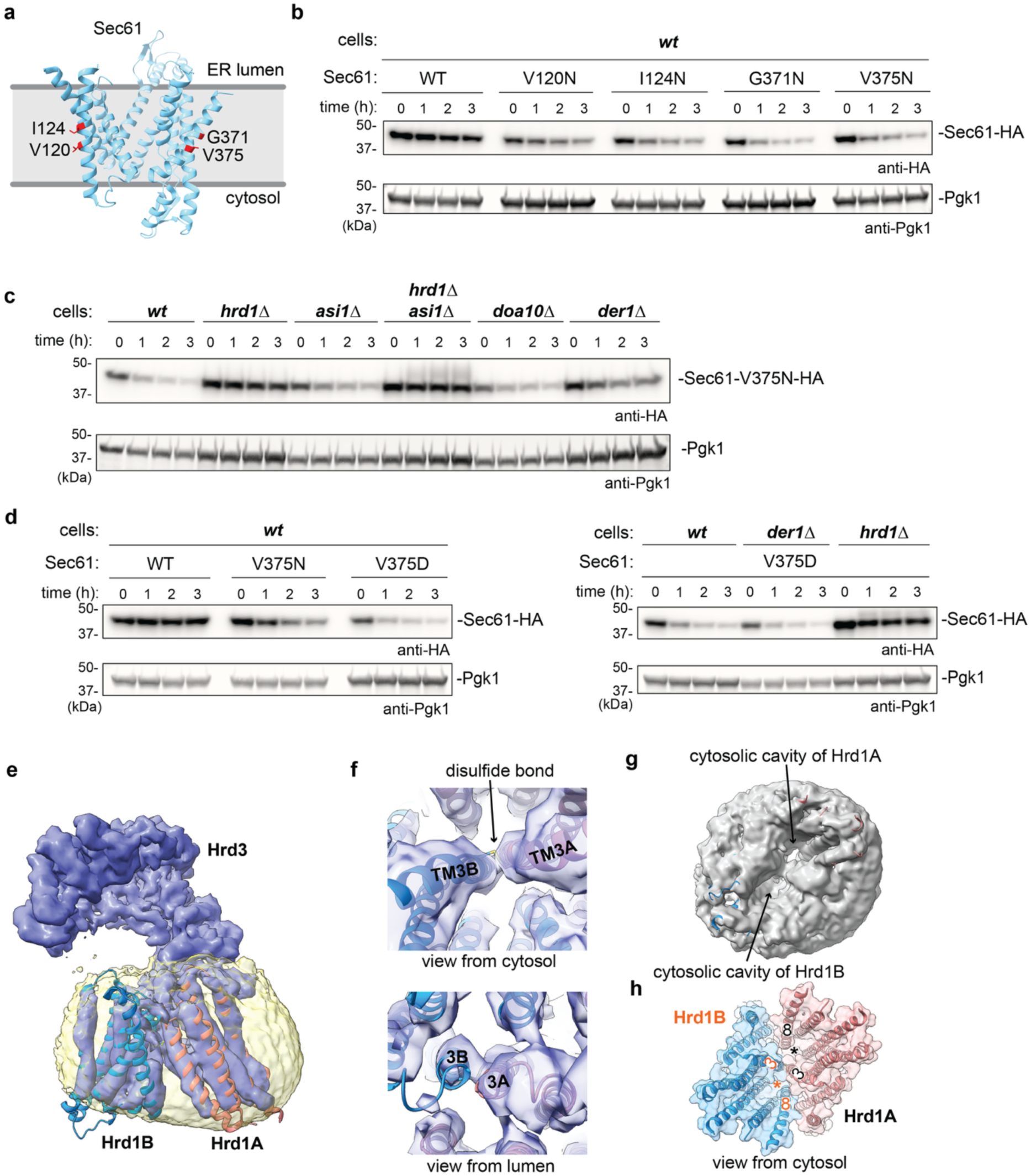
Multi-pass substrate interaction with Hrd1. (a) Asn residues were introduced into Sec61 at the indicated positions. The side chains are expected to face phospholipids inside the membrane. (b) HA-tagged wild-type (WT) Sec61 or mutants with Asn at the indicated positions were tested for degradation in wild-type (wt) cells by cycloheximide-chase experiments. The samples were analyzed by SDS-PAGE followed by immunoblotting with HA antibodies. Immunoblotting for phosphoglyceratekinase (Pgk1) served as a loading control. (c) As in (b), but for HA-Sec61 carrying the V375N mutation in cells lacking the indicated components. (d) As in (b) and (c), but for HA-Sec61 carrying either the V375N or V375D mutation. (e) CryoEM structure of a disulfide-crosslinked Hrd1 dimer complex in nanodiscs. Shown is a side view of the density map (in dark blue) with cartoon models for the two Hrd1 molecules (light blue and orange). The nanodisc density is shown in yellow. (f) Magnified views from the cytosol and lumen of the crosslinked TM3’s of the two Hrd1 molecules. (g) View of the Hrd1-containing nanodisc from the cytosol. Note the deep cavities. (h) View as in (g) with the Hrd1 molecules shown as cartoon models embedded in a semi-transparent space-filling model. The stars indicate the lateral gates formed between TMs 3 and 8.

It seemed likely that Der1 must move away from the lateral gate of Hrd1 to allow multi-pass substrates to come close to the cytosolic cavity of Hrd1. To test this idea, we used Hrd1 itself as a surrogate for how Hrd1 deals with multi-pass membrane proteins. Hrd1 is unstable even in wild-type cells ^28,29^, but particularly when overexpressed or when Hrd3 is absent ^30^. Previous experiments also showed that Der1 can readily dissociate from Hrd1 and be replaced by a second Hrd1 molecule ^12^.

To demonstrate that Hrd1 can replace Der1, we determined a cryoEM structure of an Hrd1 dimer. We introduced a Cys residue at position 91 of the lateral gate of FLAG-tagged Hrd1 (amino acids 1-326; lacking the RING finger domain) and co-expressed SBP-tagged Hrd3 (amino acids 1-767). The cells were treated with an oxidant, and the crosslinked Hrd1 dimer was purified in the detergent decylmaltose neopentyl glycol (DMNG) by binding to FLAG-antibody beads (**Extended Data Fig. 7a**). After elution, the complex was incorporated into nanodiscs, subjected to size-exclusion chromatography (SEC), and analyzed by cryoEM (**Extended Data Fig. 7b-g**) The major particle class contained two Hrd1 molecules (Hrd1A and Hrd1B) and only one Hrd3 molecule (**Fig. 6e**). Several helices in Hrd1B had weak density, likely because this Hrd1 molecule is not constrained by Hrd3. The two Hrd1 molecules are associated with one another in a nearly symmetrical fashion, but their arrangement is different from that in the inactive dimer that is likely a detergent artifact ^31^ (**Extended Data Fig. 8a** versus **8b**). Hrd1B occupies approximately the same position as Der1 in the Hrd1-Der1 structure ^11^ (**Extended Data Fig. 8c**). The TM3’s contain the crosslinked cysteines and separate the cytoplasmic cavities of the monomers (**Fig. 6f-h**). Recent cryo-EM structures of the mammalian Hrd1 dimer ^32,33^ are virtually identical to our yeast structure (**Extended Data Fig. 8d-f**), but were obtained without crosslinking, indicating that the crosslinking does not induce artifacts. Our structure shows that each Hrd1 molecule has a large cytosolic cavity (**Fig. 6g**). The structure confirms that Der1 needs to move away from Hrd1’s lateral gate for multi-pass proteins to access the thinned membrane region. It provides an explanation for why replacement of Der1 with another Hrd1 molecule causes autoubiquitination and Hrd1 degradation.

## Discussion

Our results lead to a simple and general model of ER quality control for membrane proteins. When properly folded, these proteins display hydrophobic amino acids to the surrounding lipid environment. However, when a mutation in a multi-pass protein causes its misfolding or when a subunit of a multi-component complex lacks its partner protein, hydrophilic amino acid residues may be exposed, which is energetically unfavorable. The proteins then escape the energetic penalty by partitioning into the aqueous cavity that is generated by membrane thinning next to the ubiquitin ligase Hrd1. The increased residence time near Hrd1 leads to polyubiquitination and subsequent degradation (**Fig. 7**).

**Fig. 7.**
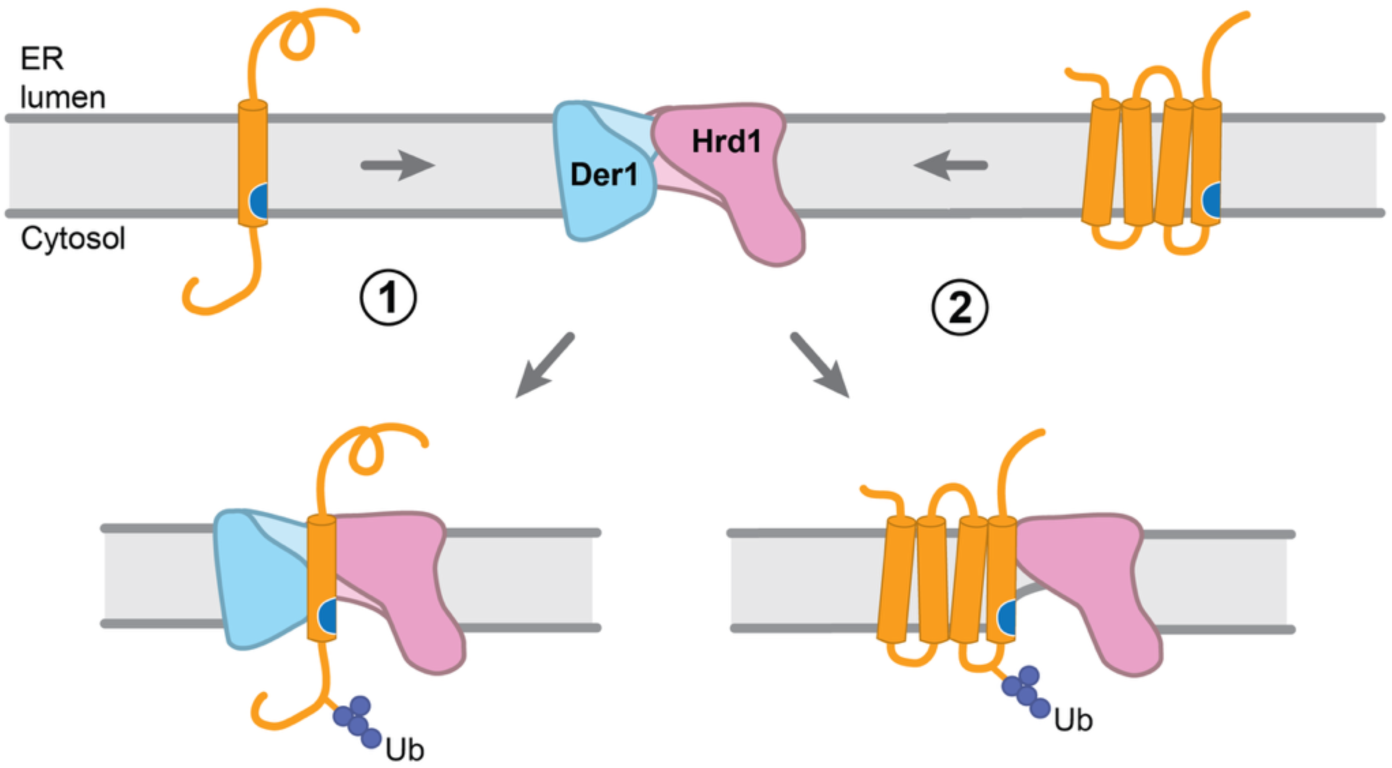
Model for the recognition of misfolded ER membrane proteins. Single-pass proteins containing a destabilizing amino acid in the TM (blue) partition to the thinned membrane region adjacent to Der1 and Hrd1 and are then polyubiquitinated (scenario 1). Multi-pass proteins partition to the cytosolic cavity of Hrd1 when Der1 is out of the way (scenario 2).

The misfolding of membrane proteins in a phospholipid bilayer is conceptually the opposite of misfolding of water-soluble proteins, as membrane proteins expose hydrophilic, rather than hydrophobic, residues. Multi-pass proteins can become unstable by mutations in cytosolic or luminal segments or by the binding of a ligand, but it seems now likely that in both cases conformational changes occur that result in the exposure of hydrophilic residues within the membrane.

Our analysis of single-pass membrane proteins shows that hydrophilic amino acids in the middle of the membrane or in the cytosolic leaflet of the lipid bilayer are most destabilizing, consistent with these residues being able to partition into the cytosolic cavity of Hrd1. Furthermore, the propensity of amino acids to trigger degradation correlates with their energetic cost in the lipid environment: the higher the cost, the more pronounced the partitioning into the aqueous cavity and the higher the rate of degradation. Specific side chain interactions may contribute, as mutations in the cytosolic cavity of Hrd1 can affect the degradation of some, but not all, membrane proteins ^34^. The partitioning model explains why degradation is relatively slow, sometimes with half-lives of more than an hour, and varies among substrates. Our results also show that mere proximity to Hrd1 does not result in substrate degradation if there is no suitable Lys residue in the cytosolic tail that can be polyubiquitinated. Surprisingly, hydrophilic TM residues in the luminal leaflet of the phospholipid bilayer do not cause degradation. For example, the D4-R construct (**Fig. 2c**) is remarkably stable, indicating that it is not well recognized by any of the ER-localized ligases. It remains to be seen whether hydrophilic residues are also more tolerated in the luminal region of multi-pass membrane proteins.

Multi-pass proteins and single-pass proteins that normally assemble with partner proteins can have hydrophilic residues in their TMs. These TMs can be inserted into the phospholipid bilayer because the energetic cost of accommodating these residues is generally compensated for by hydrophobic amino acids that favor this environment. Some TMs may be too hydrophilic to stay in the membrane on their own and could translocate into the ER lumen where they trigger ERAD-L, as demonstrated for the single-pass T-cell receptor subunits in mammalian cells ^35^. If a TM stays in the membrane and exposes hydrophilic residues to the surrounding lipids, it will ultimately partition into the thinned membrane region next to Hrd1, resulting in substrate polyubiquitination and degradation. Thus, as previously proposed ^36^, the integration of TMs into the phospholipid bilayer and their subsequent assembly are separable and consecutive events. Quality control is exerted at the assembly step and ensures that misfolded multi-pass proteins or orphaned single-pass proteins are discarded.

Interestingly, multi- and single-pass membrane proteins use different ERAD components. Previous experiments have shown that the degradation of multi-pass proteins requires Hrd1, but not Der1 ^15^. We now demonstrate that single-pass membrane proteins use both components, similarly to ERAD-L substrates. A role for Der1 had been demonstrated for several model ERAD-L substrates, but orphaned Erv25 seems to be the first substrate with wild-type sequence. The TM of single-pass proteins would enter the thinned membrane region between Der1 and Hrd1 (arrow in **Extended Data Fig. 8c**), explaining why the TM can be crosslinked to the lateral gates of both proteins (**Extended Data Fig. 5**). Consistent with most of the membrane thinning being caused by Hrd1, we found that the degradation of single-pass proteins is more dependent on Hrd1 than Der1. Der1 contributes to membrane thinning on the cytosolic side, as replacing hydrophilic with hydrophobic residues in Der1’s TM2 compromises ERAD-L ^11^. The luminal cavity of Der1 is rather shallow and probably does not facilitate the partitioning of hydrophilic TMs. Because multi-pass proteins cannot employ Der1, one might expect that they partition less efficiently than single-pass proteins to the thinned membrane region and would therefore be degraded more slowly, a prediction that is supported by our data (**Fig. 6**), but needs further testing.

In unstressed yeast cells, Hrd1 is about 5-fold more abundant than Der1 ^37^, so most Hrd1 complexes lack Der1. When the misfolding of luminal ER proteins is induced, Der1 is highly upregulated ^38^, consistent with its role in ERAD-L. However, even under these conditions, Der1 can likely dissociate from Hrd1, as the interface between the two proteins involves minimal amino acid contacts ^11,12^. Usa1 facilitates the interaction ^39,40^, but may just keep Hrd1 and Der1 in proximity while still allowing their dissociation. Hrd1 monomers may also be generated from Hrd1 dimers through autoubiquitination ^12,28,41^. Only Hrd1 monomers allow the partitioning of multi-pass membrane proteins into the thinned membrane region next to Hrd1, because substrate access is sterically blocked in Der1-Hrd1 or Hrd1-Hrd1 complexes. In addition to causing mutual polyubiquitination and degradation, Hrd1 dimers may be active in ERAD, as they generate membrane thinning comparable to Hrd1-Der1 heterodimers (**Fig. 6f** and ref.^11^). The Hrd1 dimers may mediate the basic ERAD process for luminal or single-pass proteins that is observed in the absence of all other components of the Hrd1 complex ^28,39^.

Among the three ER-localized ubiquitin ligases in *S. cerevisiae*, Hrd1 may be the only one that recognizes misfolded proteins by the proposed partitioning mechanism. The Asi ligase deals mostly with inner nuclear membrane proteins, but can also target orphan membrane proteins and at least one cytosolic protein after its transfer into the nucleus ^21,42–45^. However, the Asi complex seems to be able to weakly compete with the Hrd1 complex: all tested proteins were not affected by the deletion of *asi1* but were somewhat more stable in cells lacking both Asi1 and Hrd1 than in cells lacking only Hrd1. How exactly the Asi complex identifies its substrates remains unclear, but competition between the two ligases could determine the degree of substrate stabilization in Der1-lacking strains. The third ER-localized ubiquitin ligase, Doa10, was shown to recognize tail-anchored membrane proteins, peripheral membrane proteins, and cytosolic and nucleoplasmic proteins ^46–48^. Hydrophobic segments of these substrates seem to insert from the cytosol/nucleoplasm into a lateral, water-filled tunnel ^48^. Whereas Asi and Doa10 thus recognize their substrates by a different mechanism, the partitioning model proposed for the Hrd1 complex may apply to the Golgi, as this organelle contains the Dsc ligase complex, whose Tul1 and Dsc2 components are predicted to be structurally similar to Hrd1 and Der1, respectively ^49,50^.

To what extent our results with *S. cerevisiae* can be extended to higher eukaryotes remains to be investigated. As in yeast, the mammalian homolog of Hrd1 might recognize its substrates by partitioning to the locally thinned membrane region, and this ligase could therefore be responsible for the degradation of most misfolded ER membrane proteins. However, Hrd1 has several additional homologs, such as gp78, RNF145, and Trc8, which have distinct substrate specificities (for review, see ^8^). Presumably, the cytosolic cavities of these ligases provide more specific binding sites for substrates. Der1 has also diversified in mammals (Derlin1-3), and the homologs have different substrate specificities ^51–53^. Nevertheless, the proposed partitioning model may be the basic mechanism by which protein misfolding within membranes is recognized in all organisms. The concept may even apply to bacteria, where hydrophilic residues inside the membrane can cause degradation by FtsH ^54^, a double-pass membrane protein that contains both ATPase and protease domains. FtsH might thus combine in one polypeptide chain the three functions of ERAD components in eukaryotes: (1) recognition of a hydrophilic residue by Hrd1, (2) substrate extraction by the Cdc48 ATPase complex, and (3) substrate degradation by the proteasome.

## Methods

All chemicals were obtained from Sigma-Millipore unless specified otherwise.

### Plasmids

Yeast low-copy centromeric plasmids were derived from the pRS413 (HIS), pRS415 (LEU) or pRS416(URA) plasmid series, and 2µ high-copy plasmids were derived from pRS423, pRS425 or pRS426 ^55,56^. All constructs were assembled using Gibson assembly (NEBuilder HiFi DNA assembly Master Mix, NEB #E2621L) with custom-designed IDT gBlocks or agarose gel-purified PCR products. Vectors were linearized by restriction enzyme digests and gel purified. All PCR reactions from plasmids or gBlocks were performed using the CloneAmp polymerase (Takara #639298).

Single-pass reporter constructs were expressed from the *tef1* promoter in low-copy plasmids with or without a triple tetracycline aptamer sequence ^25^ placed upstream of the start codon. The 2A ribosome skipping sequence was derived from the ERBV1 (Equine rhinitis B virus 1) 2A peptide (GATNFSLLKLAGDVELNPGP) ^57^. The SRP-dependent signal sequence corresponds to the first 23 amino acids of S. cerevisiae Ost1 ^58^. mCherry and mNeonGreen sequences were codon optimized for expression in *S. cerevisiae* and the cysteine at position 149 in mNeonGreen was mutated to serine yielding ymoxNeonGreen. All TM sequences were flanked by GPGG…GPGG spacers. The sequence of the cytosolic tail is GPGGRVPKKNGKLSKKTN*, where RVPKKNGKLS is derived from Pdr5 C-terminal tail and the KKTN* retention motif from Wsc1 (* indicates C-terminus). Primers with scrambled codon 10 (NNK) used to create the TM10 library were synthesized by IDT.

N-glycosylation sites (<u>N</u>ST – <u>N</u> is the acceptor site) were placed into the luminal segment or the cytosolic tail, as indicated. The acceptor asparagine in the lumen was separated from the TM segment with a variable linker ^59^(15aa: <u>N</u>ST{GSGGA}SGSGGPG-TM; residues in brackets {} were eliminated in shorter linkers).

Erv25 and Emp24 genes including ∼500bp 5’ and 3’ UTRs were amplified from *S. cerevisiae* genomic DNA using PrimeStar GXL polymerase (Takara #R050A) and cloned into low-copy vectors. HA-Erv25 was constructed using PCR by placing the HA tag sequence between G20 and L21 of the native sequence. HA-Emp24 was constructed by inserting the HA sequence between A20 and H21 of the native sequence.

Plasmids for overexpression of Hrd1, Hrd3 and Der1 construct were constructed from 2µ vector backbones, with each gene placed downstream of a gal1 promoter and upstream of a cyc1 terminator.

### Cycloheximide-chase assays

Yeast cells were transformed with the indicated plasmids using the Lithium Acetate method ^60^ and grown on synthetic minimal media drop-out plates (Teknova) at 30 °C for 2 days. Fresh colonies were used to inoculate ∼4 ml liquid cultures and grown overnight at 30 °C in an incubator shaking at 230 rpm. The cultures were diluted into 50 ml of fresh media at OD_600_ ∼0.1 in 250 ml baffled Erlenmeyer flasks and grown at 30 °C, 230 rpm until the culture reached OD_600_ ∼0.5-0.8. Cells were then spun and resuspended in 10 ml media containing 100 µg/ml cycloheximide (#C7698). A 2 ml aliquot was taken (“0” min timepoint), mixed with sodium azide (0.05-0.1 %) on ice, spun in a chilled tabletop centrifuge (10,000 rpm, 30 s, 4 °C) and the pellet was flash frozen in liquid nitrogen, and stored at -80 °C. The remaining cultures were incubated at 30 °C, 230 rpm, and 2 ml-samples were taken at indicated timepoints and processed as the “0” timepoint. Cells were lysed by bead beating with equal volumes of glass beads (0.5 mm diameter, #11079105, Biospec) and lysis buffer (10 mM MOPS pH∼6.8, 1 % SDS (w/v), 8 M urea, 10 mM EDTA, 1X yeast/fungi protease inhibitor cocktail (#GB-333–1, GoldBio), 1 mM PMSF). Lysates were mixed with SDS loading buffer, heated at 65 °C for 10 min, spun (15,000 rpm, 1 min), and subjected to SDS-PAGE and immunoblotting. Signal intensities were determined in ImageStudio software (Licor) and normalized to the “0 min” timepoint. Means and standard deviations from three independent experiments were calculated in Prism (v10).

### Immunoprecipitation and Western blotting

The lysates were supplemented with SDS loading buffer, incubated at 65 °C for 10 min, spun in a table-top centrifuge (15,000 rpm, 1 min), and subjected to SDS-PAGE and immunoblotting with anti-HA antibody (clone 3F10 from rat, Roche, diluted 1:3000), anti-FLAG antibody (from rabbit, F7425, Sigma, 1:2000), anti-Pgk1 antibody (mouse monoclonal [22C5D8], #ab113687, AbCam, 1:5000), anti-mCherry polyclonal antibody (from rabbit,#26765-1-AP, Proteintech, 1:2000), anti-Myc antibody (rabbit polyclonal, #ab9106, AbCam, 1:2000), or anti-SBP antibody (mouse monoclonal, clone 20, #MAB10764, Millipore Sigma, 1:2000). Goat anti-rat HRP (ThermoFisher, #31470), IRDye goat anti-mouse-680RD (#926–68070, Licor) and IRDye donkey anti-rabbit-800CW (#926-32213, Licor) secondary antibodies were and used for visualization using Amersham ImageQuant or LicorM imagers. Signal intensities were determined in ImageStudio software (Licor)and normalized to the “0” timepoint. Means and standard deviations from three independent experiments were calculated in Prism (v10).

### Flow cytometry

Yeast cells were transformed with the indicated plasmids and grown to OD_600_∼0.4-0.6 in liquid SC drop-out media. The samples were filtered through a 35µm mesh strainer and used immediately. Data were collected on Becto Dickinson LSR II flow cytometer or on an Attune NxT flow cytometer and data analysis was performed using FlowJo. Samples were gated for live and single cells. All flow cytometry experiments are representative of at least two independent replicates.

### Cell sorting

Yeast cells transformed with the TM10 library were grown to OD_600_ ∼0.4-0.6 and mixed with 500 µM tetracycline for 1 h before sorting on a Sony SH800 cell sorter using a 100 µm chip. Cells expressing mCherry and varying levels of mNeonGreen (mNG) were sorted into two bins (mNG high and mNG low, ∼300,000 cells per bin) and immediately mixed with 2x SC-URA media. The collected cells were grown overnight, spun, and plasmids were extracted using the yeast plasmid extraction kit (ZymoResearch #D2004). Extracted plasmids were used as a template for PCR (CloneAmp polymerase, Takara #639298) to amplify a fragment containing the TM sequence using primers flanked with Illumina adaptors. PCR reactions were purified using the QIAquick PCR purification kit (Qiagen #28104), and confirmed to yield a single (∼350bp) fragment by agarose gel electrophoresis. The samples were analyzed by the AmpExp sequencing service provided by Quintara Co Ltd. Read counts from abundance analysis were used to calculate percentages of amino acid enrichment in sorted bins over the pre-sorted library using R software. Relative abundances in each bin (mNG-high and mNG-low) were determined by normalizing the percentage of amino acid enrichment in each bin by the percentage of the same amino acid in a pre-sorted population. Truncated and misaligned reads were excluded from analyses. Log_2_TMSS (log_2_ TM Stability Score) was calculated as log_2_ ratio of pseudocount-adjusted relative abundances in mNG-high and mNG-low bins: log_2_((mNG-high + 10^-6^)/(mNG-low + 10^-6^)).

### Disulfide crosslinking

The indicated strains were transformed with plasmids and selected on synthetic medium drop-out plates (Teknova) for two days at 30 °C. Multiple colonies were picked to inoculate a starter culture (4 ml). After overnight incubation at 30 °C, the cells were diluted to 50–100 ml at OD_600_∼ 0.1 and grown at 30 °C until the OD_600_ reached 0.5 to 0.8. The cells were then spun at 3000 rpm for 3 min at room temperature (RT), and the cell pellet resuspended in 5 ml of pre-warmed (30 °C) PBS or SC minimal media supplemented with 500 μM 4,4’-dithiopyridine (Aldrithiol-4, #143057). The reactions we incubated for 30 min at 30 °C in a shaker (230 rpm), followed by a quench reaction with 10 mM N-ethylmaleimide (#E3876) for 15 min on ice. Cells were spun (10,000 rpm, 4 °C, 1 min), flash frozen in liquid nitrogen, and stored at -80 °C before use.

### Protein overexpression and purification

Hrd1(1-326)FLAG-I91C and Hrd3(1-767)tevSBP were co-transformed into a *ubc7 -/-* knockout diploid strain (Horizon Discovery) and grown on selective minimal media plates for two days. Multiple colonies were picked to inoculate a 50 ml pre-culture. After growth overnight at 30 °C, the culture was used to inoculate 6 x 0.6 L fresh minimal media in 2 L baffled flasks. The cultures were grown for 24-30 hours at 30 °C and protein expression was induced by addition of 2 % galactose in minimal media supplemented with glutamine as additional nitrogen source (5 g/L). The cells were shifted to 25 °C and grown for additional 12-14 hours. The cultures were then spun (3000 rpm, 5 min) and the pellets immediately resuspended in 400 ml of media or PBS buffer prewarmed to 30 °C, supplemented with 500 µM 4,4’-dithiopyridine, and shaken for 30 min at 30 °C. The reactions were quenched with 1 mM N-ethylmaleimide on ice followed by spinning the cells and flash-freezing the pellets in liquid nitrogen. The cells were lysed by bead beating (2 s ON, 40 s OFF cycle, 40-60 min, Biospec bead beater) with glass beads (0.5 mm diameter, #11079105, Biospec) in buffer A (50 mM HEPES.NaOH pH7.5, 400 mM NaCl) supplemented with 1 mM PMSF and 1x yeast fungi protease inhibitor cocktail (lysis buffer), and the lysates were spun for 10 min (3000 rpm, 4 °C) to sediment unlysed cells and debris. The supernatant was transferred to Ti45 tubes and membranes were sedimented at 50,000 rpm for 1 h at 4 °C in a Beckman ultracentrifuge. Pelleted membranes were washed and dounced in lysis buffer and spun again before flash freezing in liquid nitrogen. Membranes were solubilized on a rotator with buffer A supplemented with 1 % DMNG for 1 h at 4 °C and then spun (50,000 rpm, 4 °C, 30 min) to sediment insoluble material. The cleared supernatant was mixed with anti-FLAG M2 agarose resin for 1.5h, washed with buffer B (25 mM HEPES.NaOH pH7.5, 400 mM NaCl) supplemented with 0.1 % DMNG. Bound proteins were eluted with buffer C (25 mM HEPES.NaOH pH7.5, 250 mM NaCl) supplemented with 0.1 % digitonin, 0.1 mg/ml 3xFLAG peptide and 10 % glycerol. Eluted protein was concentrated using a 100 kDa MWCO Amicon device and flash-frozen or used immediately.

### Reconstitution into nanodisc

For nanodisc reconstitution, the protein was mixed with yeast polar lipids (#190001, Avanti Polar Lipids). The lipid stock was prepared as in ^61^. Briefly, the chloroform lipid mixture was first dried under a nitrogen gas stream and then under vacuum overnight to yield a thin lipid film in a glass vial. The lipid film was then resuspended in buffer (25 mM HEPES, pH 7.4, 200 mM NaCl) in a water bath sonicator (Brandon) and subjected to 10 freeze-thaw cycles in liquid nitrogen and water sonication bath at room temperature. The lipid stock (10 mM) was then used directly for nanodisc reconstitution or flash frozen in liquid nitrogen for later use.

Proteins purified in digitonin were incorporated into nanodisc using a 1:240:3 molar ratio of protein:lipid:membrane-scaffold protein. Commercial MSP1D1 (#M6574) was used for reconstitution of the Hrd1-Hrd1 dimer complex. The mixture was incubated at 4 °C for 1 h while rotating. Bio-Beads (Bio-Rad) were then added, and the sample was slowly rotated at 4 °C overnight. The Bio-Beads were removed, and the mixture was centrifuged at 12,000g for 10 min to remove aggregates and sediment lipid vesicles. The supernatant was loaded onto a Superose 6 3.2/300 Increase size-exclusion column (GE Healthcare) equilibrated in gel-filtration buffer (25 mM HEPES pH 7.4 and 200 mM NaCl). Fractions containing nanodisc-reconstituted complexes were pooled and concentrated with a 100kDa MWCO device (Amicon).

### Single-particle cryo-EM sample preparation and data acquisition

A total of 3 µl of the concentrated sample was applied to glow-discharged (Pelco easiGlow, 15 mA, glow 45 s, hold 15 s, 0.38 mBar) grids (Quantifoil R1.2/1.3, Au 300 mesh). The grids were blotted for 5-7 s at approximately 95%-100% humidity and plunge-frozen in liquid ethane using a Leica GP2 plunger instrument. The grids were screened on a Glacios-2 (Thermo) electron microscope operated with Epu software (Thermo) at 200 kV. The microscope was equipped with X-FEG electron source, Selectris energy filter, and Falcon 4i camera. For the Hrd1-Hrd1 dimer sample, a larger dataset was collected on a Titan Krios electron microscope (FEI) operated at 300 kV and equipped with a high-brightness field emission gun (X-FEG), K3 direct electron detector (Gatan), spherical aberration corrector (Cs ∼0.01), and a Gatan BioContinuum HD energy filter (slit width 8eV) at the HHMI Janelia Farm Cryo-EM facility (Krios 1).

Cryo-EM movies for the Hrd1-Hrd1 dimer sample were recorded in super-resolution counting mode using SerialEM. The nominal magnification of ×81,000 corresponds to a calibrated physical pixel size of 0.857 Å and 0.4285 Å in the super-resolution mode. The total dose was 55 electrons Å^−2^ spread over 50 frame movie (dose per frame 1.1 electrons per Å^2^). The defocus was between −0.8 and −2.0 µm.

### Data processing

The image processing workflows are illustrated in detail in **Extended Data Fig. 7** and were performed using cryoSPARC v4.4-4.7. Briefly, motion-correction and dose weighting were performed using the Patch_Motion_Correction module in cryoSPARC ^62^. Next, CTF was estimated with the Patch_CTF module and micrographs with a CTF-estimated resolution worse than 6 Å, extreme defocus range, or high drift values were excluded from further data analysis. Initial particles were picked using cryoSPARC’s Blob picker module. Two-dimensional (2D) classifications were performed and 2D averages showing expected protein features were used as initial templates for particle picking across all selected micrographs using the Template_Picker module. After another round of 2D classification, particles belonging to acceptable 2D class averages were selected for an Ab-initio 3D reconstruction. Secondary structure features of the Hrd3 luminal domain were readily visible in the initial 3D reconstruction. This initial 3D volume was then utilized to generate 50 2D templates using the Create_Templates module. These 2D templates served as templates for particle picking across all selected micrographs again. After 2D classification, selected particles were subjected to another round of Ab-initio reconstruction. Particles from good ab-initio volumes were used for 3D classification. 3D classes with two Hrd1 molecules were selected for Non-Uniform refinement^63^. Local resolution variations were estimated in cryoSPARC. Resolutions of the refinement were estimated according to the gold-standard Fourier Shell Correlation (FSC) 0.143 criterion. Software and servers used in this project were supported by SBGrid (http://sbgrid.org) ^64^ and the HMS Research Computing initiative.

### Model building and refinement

The initial Hrd1-Hrd3 model (from pdb ID: 6VJY) and the second copy of Hrd1 (from 6VJY) were first docked into the cryo-EM density map using UCSF ChimeraX ^65^. The model was manually adjusted in Coot ^66,67^ and refined using Phenix ^68^. Figures were prepared using USCF ChimeraX 1.9-1.10 and Adobe Illustrator (v26.5).

### Fluorescence microscopy

BY4741 cells transformed with the indicated plasmids and grown in drop-out minimal media (low-fluorescence YNB, Formedium #CYN6505) to OD600∼0.5 at 30°C before imaging. The cells were immobilized on glass coverslip coated with concanavalin A and imaged immediately using a confocal fluorescence microscope (Nikon Ti inverted microscope with Prior Proscan II motorized stage and shutters, a 16-bit Hamamatsu ORCA-Fusion BT sCMOS camera, Nikon LUN-F XL solid state lasers (405, 488, 561, and 640 wavelengths), and 40X, 63X, and 100X Plan Apo 1.4 or 1.44 N/A oil objectives). All images were acquired using the 100X Plan Apo 1.44 NA oil objective using NIS-Elements (v5.21.03) software. Image processing and figure preparation was performed in FiJi (ImageJ2 v 2.9.0/1.53t) and Adobe Illustrator (v26.5).

## Data Availability Statement

All data are available in the main text, extended data, or the supplementary materials. The cryo-EM map of the Hrd1-Hrd1 dimer was deposited in the Electron Microscopy Data Bank (EMDB-73699). The model coordinates were deposited in the Protein Data Bank (PDB ID 9Z0E). Reagents generated by this study are available from the corresponding author with a completed materials transfer agreement.

## Code Availability Statement

No custom computer code, algorithm, or software was generated.

## Acknowledgments

We thank Susan Shao, Ryan Baldridge, and Yoko Shibata for critical reading of the manuscript. We are thankful to Richard Walsh at the Harvard Cryo-EM Center for Structural Biology for help with microscope operation; Chad Hicks at the BU Cryo-EM center for help with microscope operation and data collection; Rui Yan, Nicholas Spellman, and Shixin Yang for their help with data collection at the cryo-EM facility at the HHMI Janelia Research Campus; Shaun Rawson for help with data processing; SBGrid team for software and workstation support. We thank the Core for Imaging and Education (CITE) at Harvard Medical School for technical assistance. This work was funded by National Institutes of Health grant R01GMO52586 (T.A.R.), the Howard Hughes Medical Institute (HHMI) (T.A.R.), the American Heart Association grant #833683/2021 (R.P.), the Charles A. King Trust Postdoctoral Research Fellowship Program, Bank of America, N.A., Co-Trustees (R.P.), and the 1K99GM154114-01 grant from NIH/NIGMS (R.P.). This manuscript is the result of funding in whole or in part by the National Institutes of Health (NIH). It is subject to the NIH Public Access Policy. Through acceptance of this federal funding, NIH has been given a right to make this manuscript publicly available in PubMed Central upon the Official Date of Publication, as defined by NIH.

## Author contributions

R.P. performed all experiments and analyzed data. R.P. and T.A.R conceptualized the study, designed experiments, obtained funding, and wrote the manuscript.

---

**Extended Data Fig. 1.**
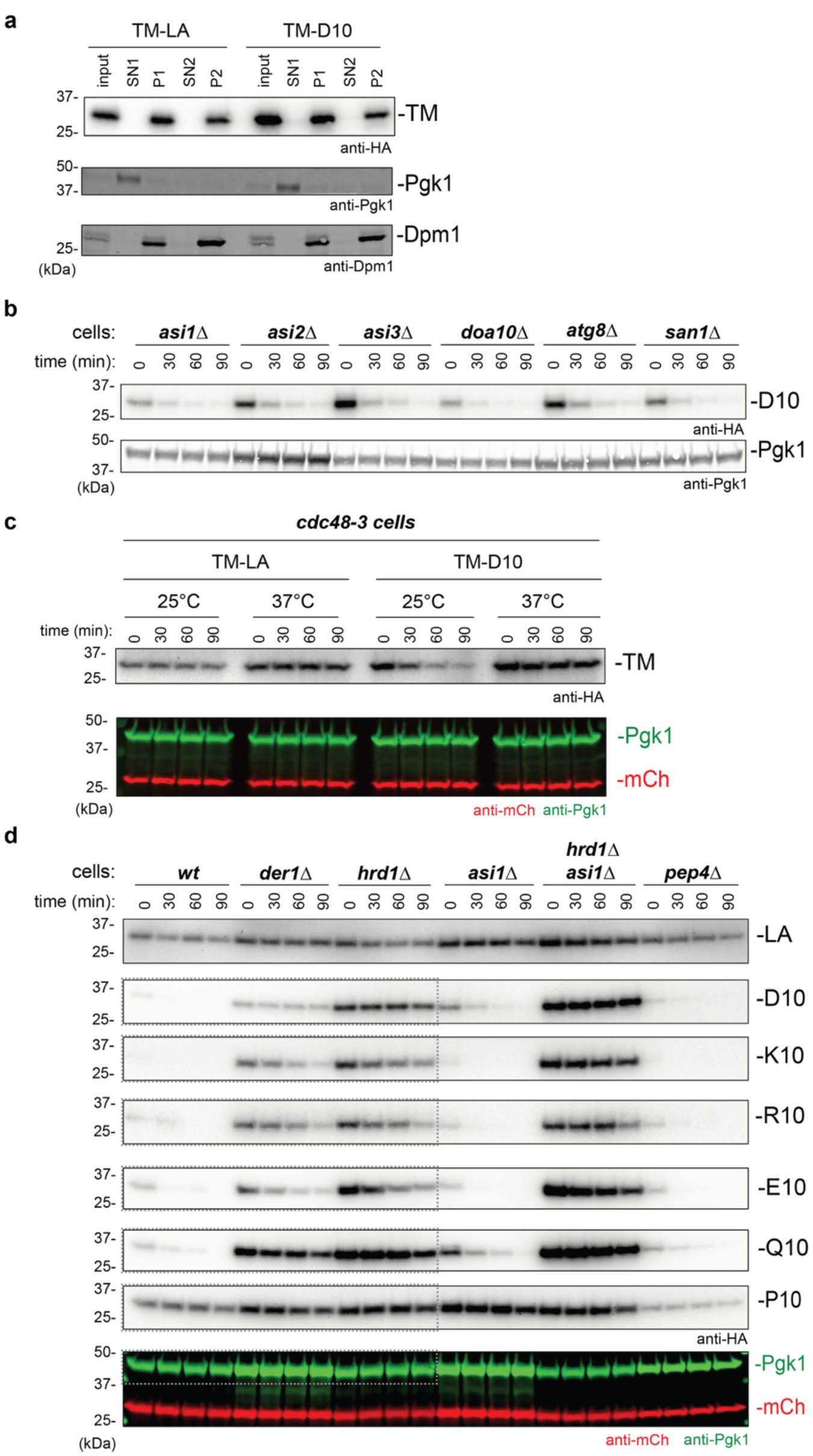
Hydrophilic amino acids in an idealized TM trigger ERAD. (a) The membrane localization of HA-tagged LA and D10 constructs was tested by cell fractionation. Both constructs were expressed in Hrd1-lacking cells to prevent their degradation. Cells were either analyzed directly (input) or fractionated into a supernatant (SN1) and membrane pellet (P1). The membrane pellet was resuspended and again subjected to centrifugation to yield a supernatant (SN2) and pellet (P2). All samples were subjected to SDS-PAGE and immunoblotting with anti-HA antibodies. Immunoblotting for phosphoglyceratekinase (Pgk1) and dolichol-phosphate mannosyltransferase (Dpm1) served as cytosolic and ER membrane markers, respectively. (b) The HA-tagged D10 construct was expressed in cells lacking the indicated components. The degradation of D10 was followed by cycloheximide-chase experiments. The samples were analyzed by SDS-PAGE and immunoblotting with anti-HA antibodies. Immunoblotting for phosphoglyceratekinase (Pgk1) served as a loading control. (c) The HA-tagged LA and D10 constructs were expressed in *cdc48-3* cells that contain a temperature-sensitive Cdc48 mutation. The cells were incubated at either the permissive (25 °C) or non-permissive (37 °C) temperature and subjected to cycloheximide-chase experiments. The samples were analyzed as in (b). Immunoblotting for phosphoglyceratekinase (Pgk1) and mCh served as loading controls. (d) Constructs with the hydrophobic TM (LA) or with a TM containing the indicated amino acids at position 10 were expressed in wt cells or cells lacking the indicated components. Cycloheximide-chase experiments were performed and the samples analyzed as in (c). Dotted lines indicate data shown in Fig. 1e; the loading control for the P10 construct is shown.

**Extended Data Fig. 2.**
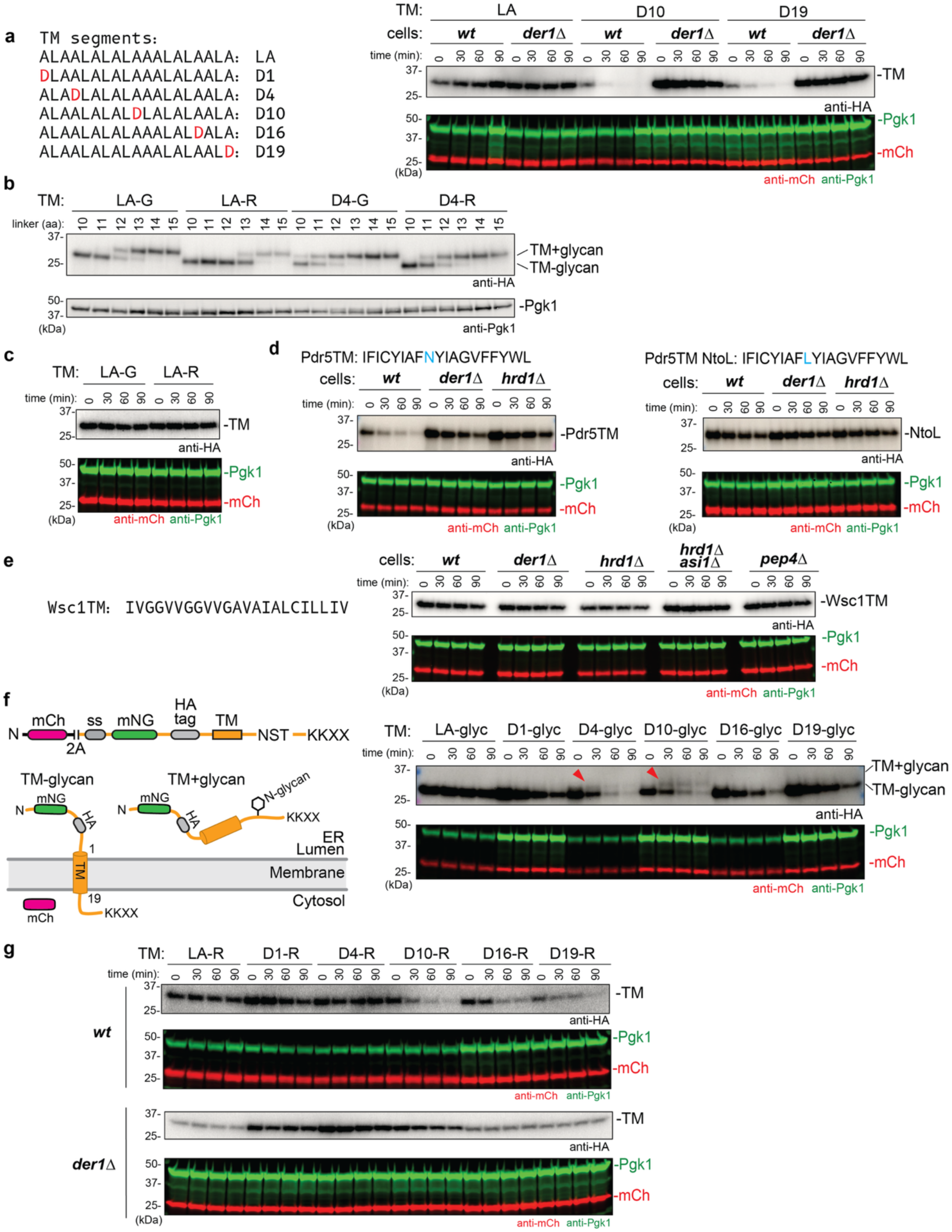
Hydrophilic TMs cause ERAD without being translocated into the ER lumen. (a) HA-tagged constructs containing the hydrophobic LA segment or TMs with Asp residues at the indicated positions were tested in wild-type (wt) cells or cells lacking Der1 for their degradation by cycloheximide-chase experiments. The samples were subjected to SDS-PAGE and immunoblotting with anti-HA antibodies. Immunoblotting for phosphoglyceratekinase (Pgk1) and mCh served as loading controls. (b) HA-tagged LA or D4 constructs with GPGG (-G) or RPRR (-R) linkers at the cytosolic end of the TM and N-glycosylation sites at different luminal positions were expressed in cells lacking Hrd1 and Asi1. The attachment of a glycan was determined by the mobility shift in anti-HA antibody immunoblots. Immunoblotting for phosphoglyceratekinase (Pgk1) served as a loading control. (c) The degradation of LA constructs with GPGG (-G) or RPRR (-R) linkers at the cytosolic end of the TM was tested by cycloheximide-chase experiments as in (a). (d) A TM of the multi-pass membrane protein Pdr5 was placed into the reporter construct. The TM contains an Asn residue (in blue) that was mutated to Leu for the experiments on the right. The constructs were expressed in wild-type (wt) cells or cells lacking the indicated components, and cycloheximide-chase experiments were performed as in (a). (e) As in (d), but with the entirely hydrophobic TM of the Wsc1 protein. (f) To test whether the TMs serve as a stop-transfer sequences, a N-glycosylation site was introduced into the cytosolic tail of the LA construct or into constructs containing Asp residues at the indicated positions (the glycosylation site NST is 10 residues downstream of the last TM residue). Cycloheximide-chase experiments were performed as in (a). Glycosylated species have an increased molecular weight (red arrowheads). (g) The RPRR linker was placed at the cytosolic end of the indicated TMs to prevent sliding of the TM towards the ER lumen. Cycloheximide-chase experiments were performed with wt cells or cells lacking Der1 as in (a).

**Extended Data Fig. 3.**
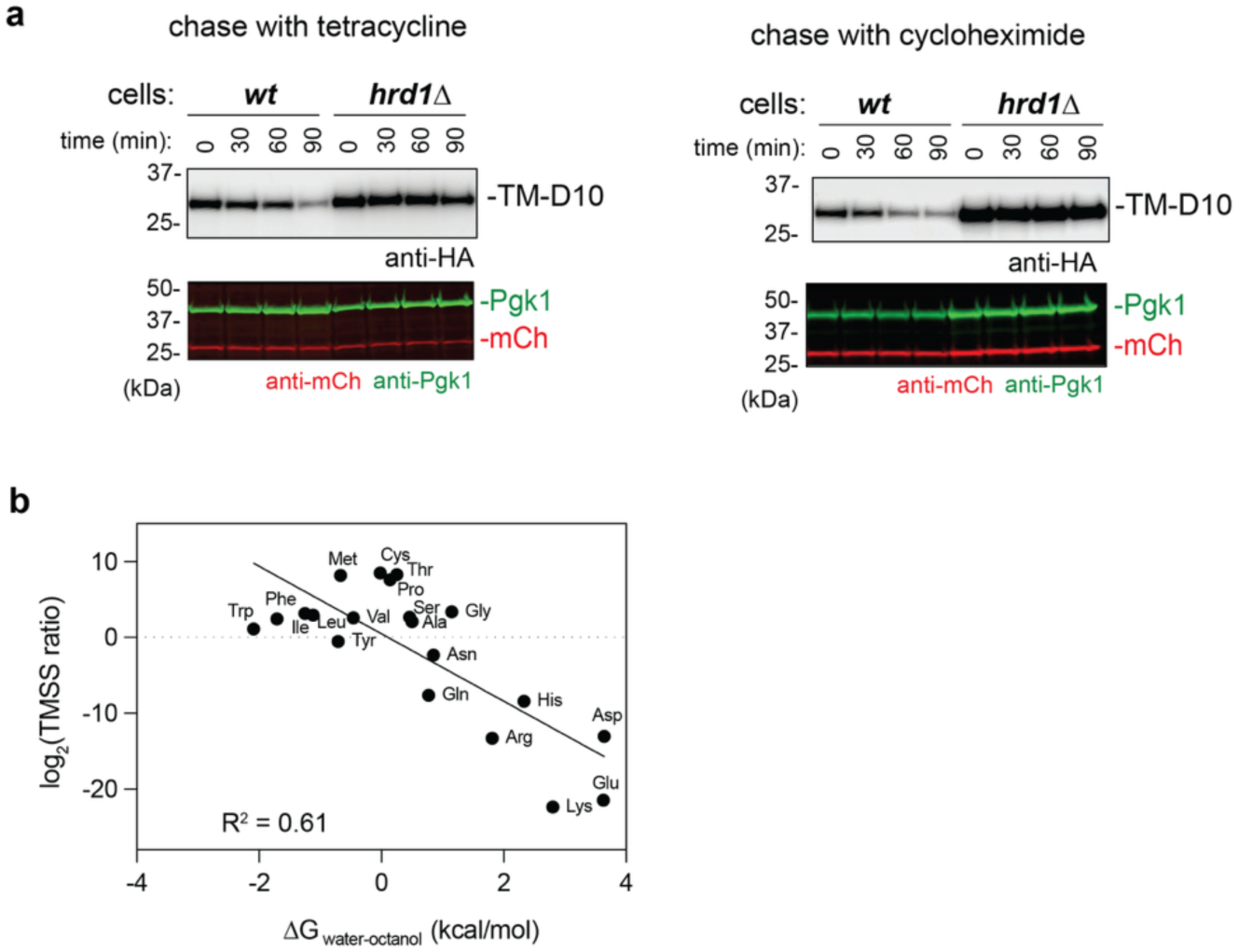
Correlation between the destabilizing effect of an amino acid and its propensity to partition into octanol. (a) Left: The HA-tagged D10 construct was placed under the control of the *tef1* promoter and tetracycline aptamers to allow selective shut-off of expression. The construct was expressed in wild-type (wt) cells or cells lacking Hrd1. After addition of tetracyclin, samples were taken at different time points and analyzed by SDS-PAGE and immunoblotting with anti-HA antibodies. Immunoblotting for phosphoglyceratekinase (Pgk1) and mCh served as loading controls. Right: The degradation of D10 was followed after addition of cycloheximide to stop the expression of all proteins. (b) The TM stability score (TMSS), calculated as the ratio of the relative depletion in the high mNG/mCh population over the enrichment in the low mNG/mCh population, was plotted against ΔG_water-octanol_, the free energy required to move an amino acid from water into octanol^17^.

**Extended Data Fig. 4.**
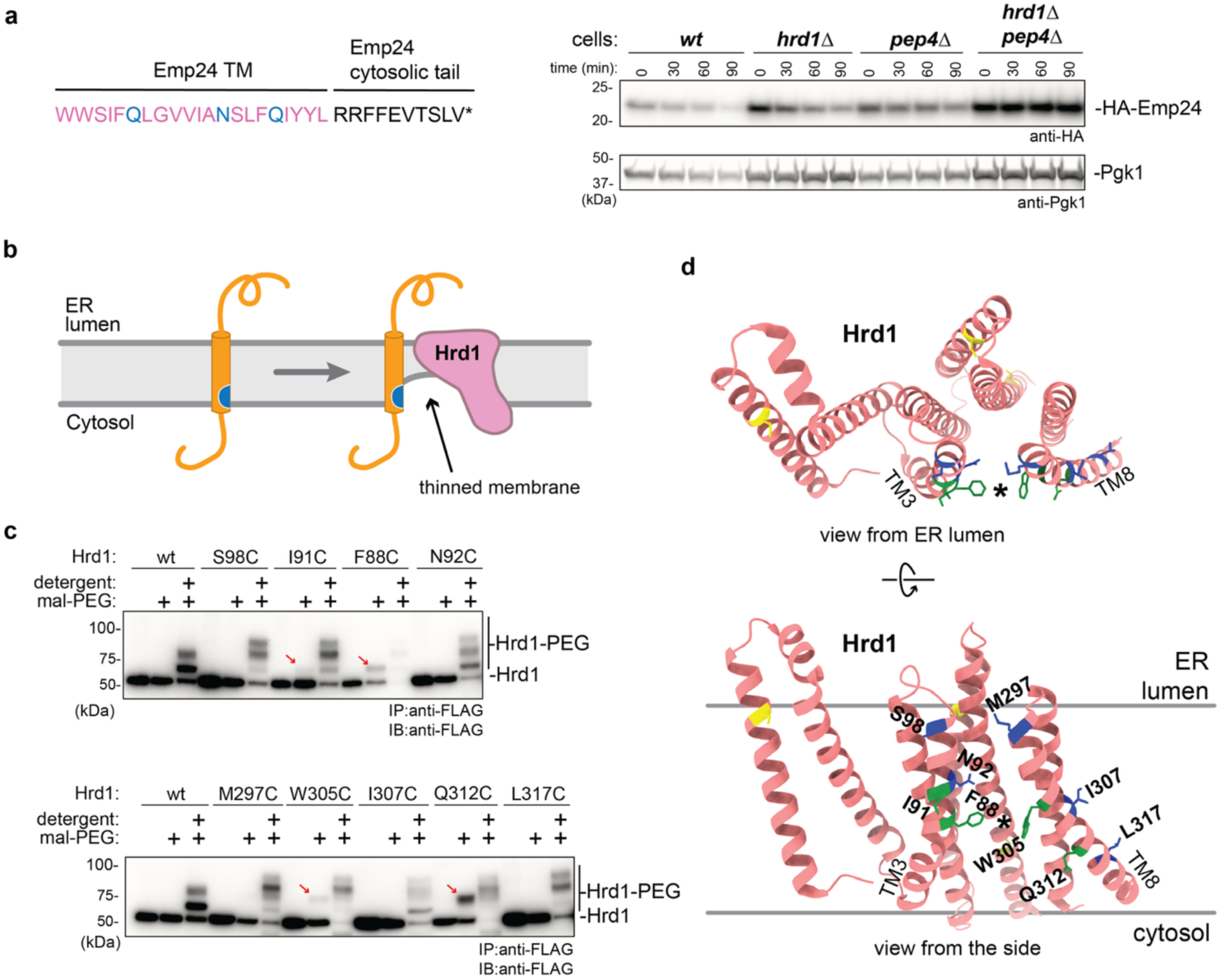
Hrd1 has a cytosolic cavity. (a) HA-Emp24 was expressed from a CEN plasmid under the endogenous promoter in wild-type (wt) cells or cells lacking the indicated components. Cycloheximide-chase experiments were performed and the samples analyzed by SDS-PAGE and blotting with anti-HA antibodies. Immunoblotting for phosphoglyceratekinase (Pgk1) served as loading control. (b) Model for the partitioning of a TM with a hydrophilic residue (in blue) into the aqueous environment of Hrd1’s cytosolic cavity. (c) Membranes from cells containing FLAG-tagged wild-type (wt) Hrd1 or Hrd1 with Cys residues introduced at different positions were incubated with 5 kDa maleimide-polyethyleneglycol (mal-PEG) in the absence or presence of detergent, as indicated. Unreacted mal-PEG was quenched with cysteine followed by detergent solubilization and immunoprecipitation with anti-FLAG beads. The samples were analyzed by SDS-PAGE and blotting with anti-FLAG antibodies. The red arrows point to Hrd1 modified by mal-PEG in the absence of detergent. (d) Residues tested for mal-PEG modification are shown in a cartoon model of Hrd1 (pdbID: 6VJY), viewed from the ER lumen (top panel) and from the side (lower panel). Residues modified by mal-PEG are highlighted in green, residues resistant to modification in blue, and endogenous cysteines in yellow. The lateral gate is indicated by an asterisk.

**Extended Data Fig. 5.**
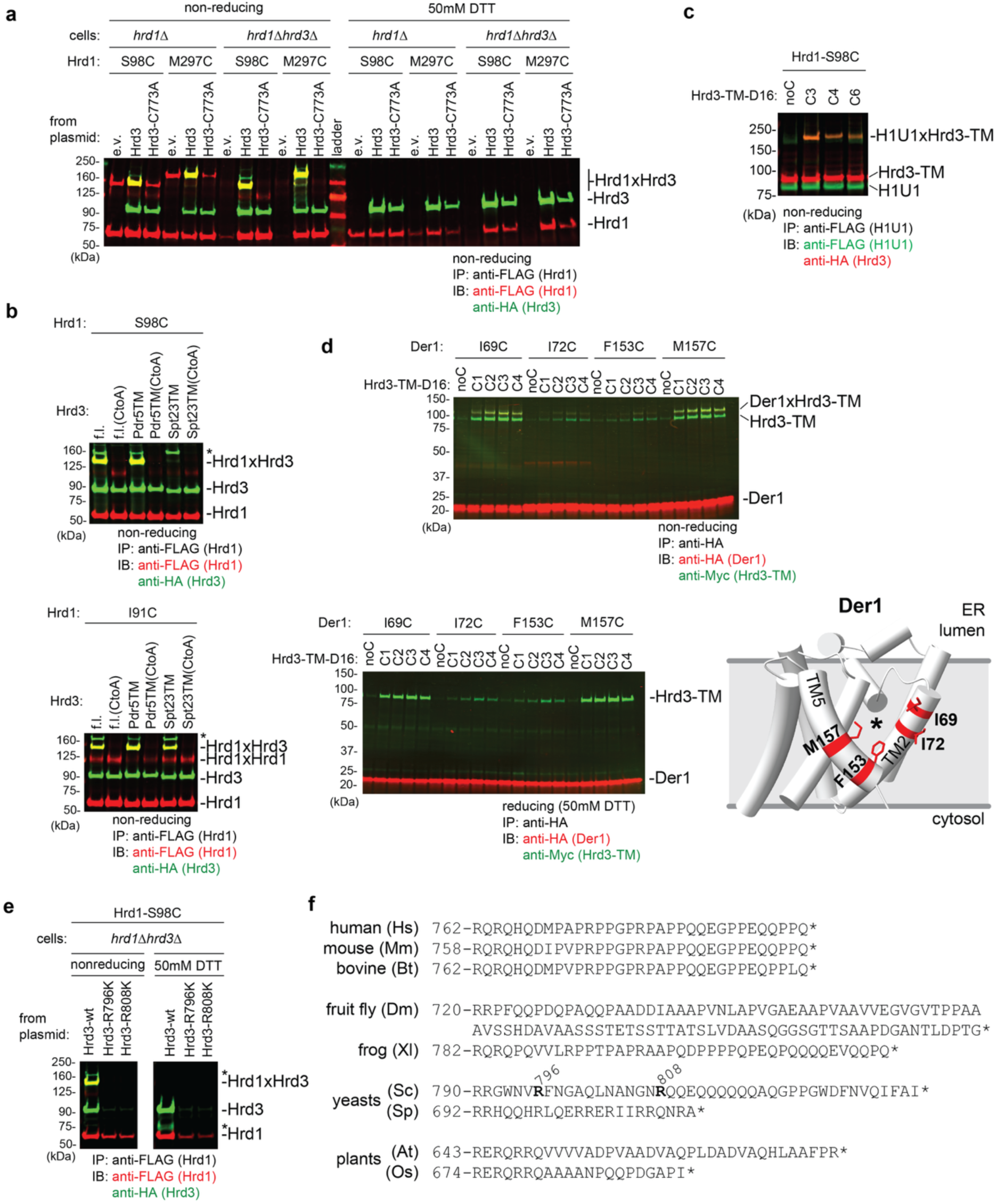
Disulfide crosslinking of Hrd3’s TM to the lateral gates of Hrd1 and Der1. (a) Disulfide crosslinking of the TM of Hrd3 with the lateral gate of Hrd1. HA-tagged Hrd3 with endogenous Cys773 in the TM (LVTMG**C**ILGIFLLSILMSTLAA) was expressed together with FLAG-tagged Hrd1 containing cysteines at the indicated positions in cells lacking Hrd1 or both Hrd1 and Hrd3. Where indicated, an empty vector (e.v.) was used or Cys773 in Hrd3’s TM was mutated to Ala. Disulfide bond formation was induced with an oxidant, and a detergent extract subjected to immunoprecipitation with anti-FLAG beads. The samples were analyzed by non-reducing or reducing (50 mM DTT) SDS-PAGE, followed by blotting with anti-HA and anti-FLAG antibodies. (b) As in (a), but where indicated, the TM of full-length (f.l.) Hrd3 was replaced by TMs from Pdr5 or Spt23. Hrd1 contained a Cys at position 98 (upper panel) or 91 (lower panel). Controls were performed with mutants in which the Cys in the TMs was mutated to Ala (CtoA). (c) As in (a), but with Hrd3’s TM replaced with an idealized TM containing D16 and Cys at the indicated positions. (d) Disulfide crosslinking of the TM of Hrd3 with the lateral gate of Der1. Myc-tagged Hrd3 with an idealized TM containing D16 and Cys at various positions was expressed together with HA-tagged Der1 containing Cys at the indicated positions in cells lacking both Hrd3 and Der1. Disulfide bond formation was induced with an oxidant, and a detergent extract subjected to immunoprecipitation with anti-HA beads. The samples were analyzed by non-reducing SDS-PAGE, followed by blotting with anti-HA and anti-Myc antibodies (upper panel). In the lower panel, the samples were reduced with 50 mM DTT before SDS-PAGE. The lower right panel shows a cartoon of the Der1 structure with crosslinking residues highlighted in red. (e) HA-tagged wild-type (wt) Hrd3 or Hrd3 containing a Lys residue at two different positions was expressed together with FLAG-tagged Hrd1 containing a Cys at position 98 in the lateral gate. Disulfide bond formation was induced with an oxidant, and a detergent extract subjected to immunoprecipitation with anti-FLAG beads. The samples were analyzed by non-reducing or reducing (50 mM DTT) SDS-PAGE, followed by blotting with anti-HA and anti-FLAG antibodies. The stars point to unidentified crosslinked products. (f) Sequences of the cytosolic tails of Hrd3 from different species. The star indicates the C-terminus. Hs, *Homo sapiens*; Mm, *Mus musculus*; Bt, *Bos taurus*; Dm, *Drosophila melanogaster*; Xl, *Xenopus laevis*; Sc, *Saccharomyces cerevisiae*; Sp*, Schizosaccharomyces pombe*; At, *Arabidopsis thaliana*; Os, *Oryza sativa*. Note the absence of Lys residues.

**Extended Data Fig. 6.**
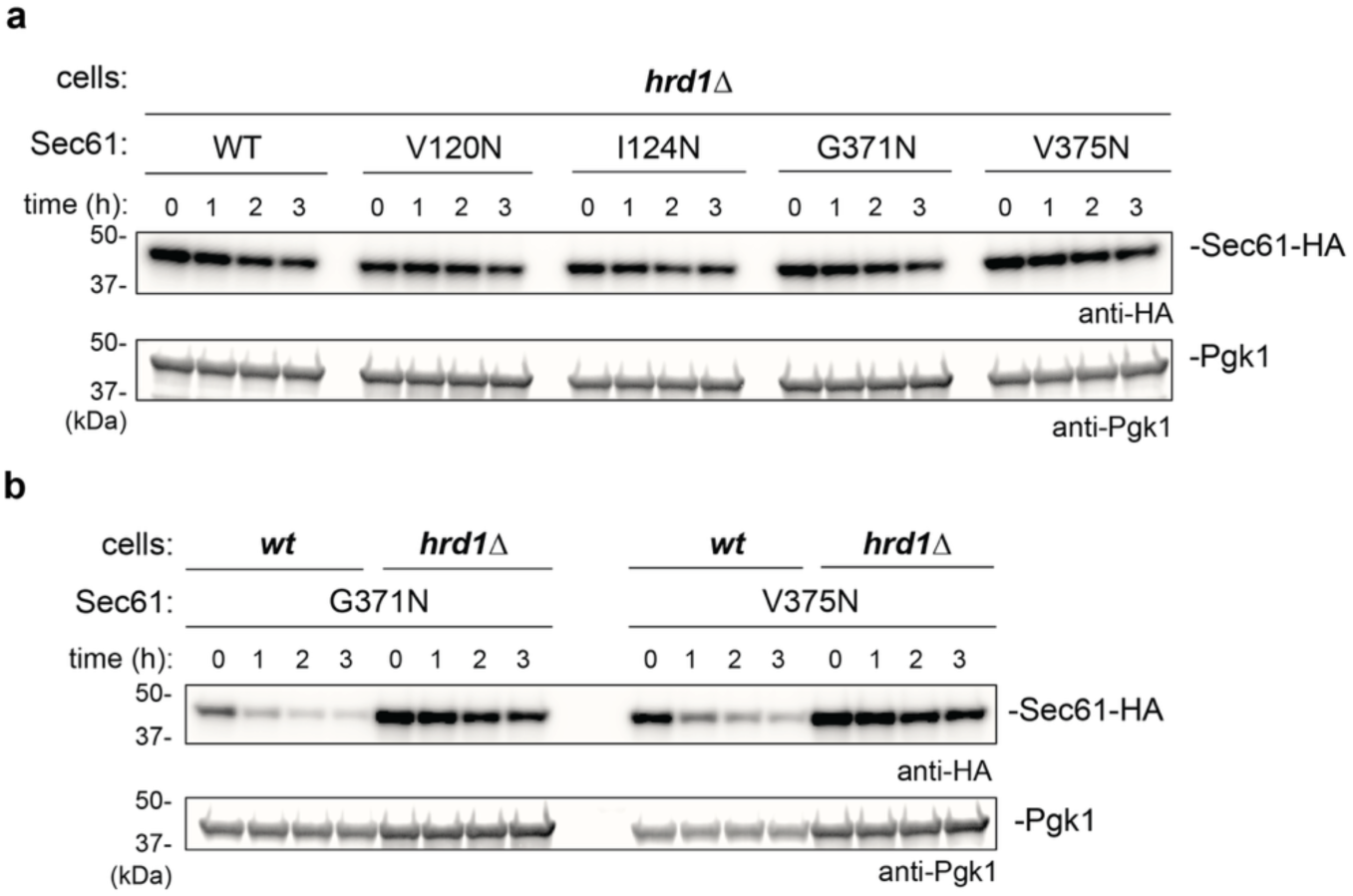
Sec61 mutants are degraded in a Hrd1-dependent and Der1-independent manner. (a) HA-tagged wild-type (WT) Sec61 or mutants with Asn at the indicated positions were tested for degradation in cells lacking Hrd1 by cycloheximide-chase experiments. The samples were analyzed by SDS-PAGE followed by immunoblotting with anti-HA antibodies. Immunoblotting for phosphoglyceratekinase (Pgk1) served as a loading control. Note that the mutants are stable, whereas they are degraded in wild-type cells (Fig. 6b). (b) HA-Sec61 with position 375 mutated to either Asn or Asp was tested for degradation in wild-type (wt) cells or cells lacking the indicated components.

**Extended Data Fig. 7.**
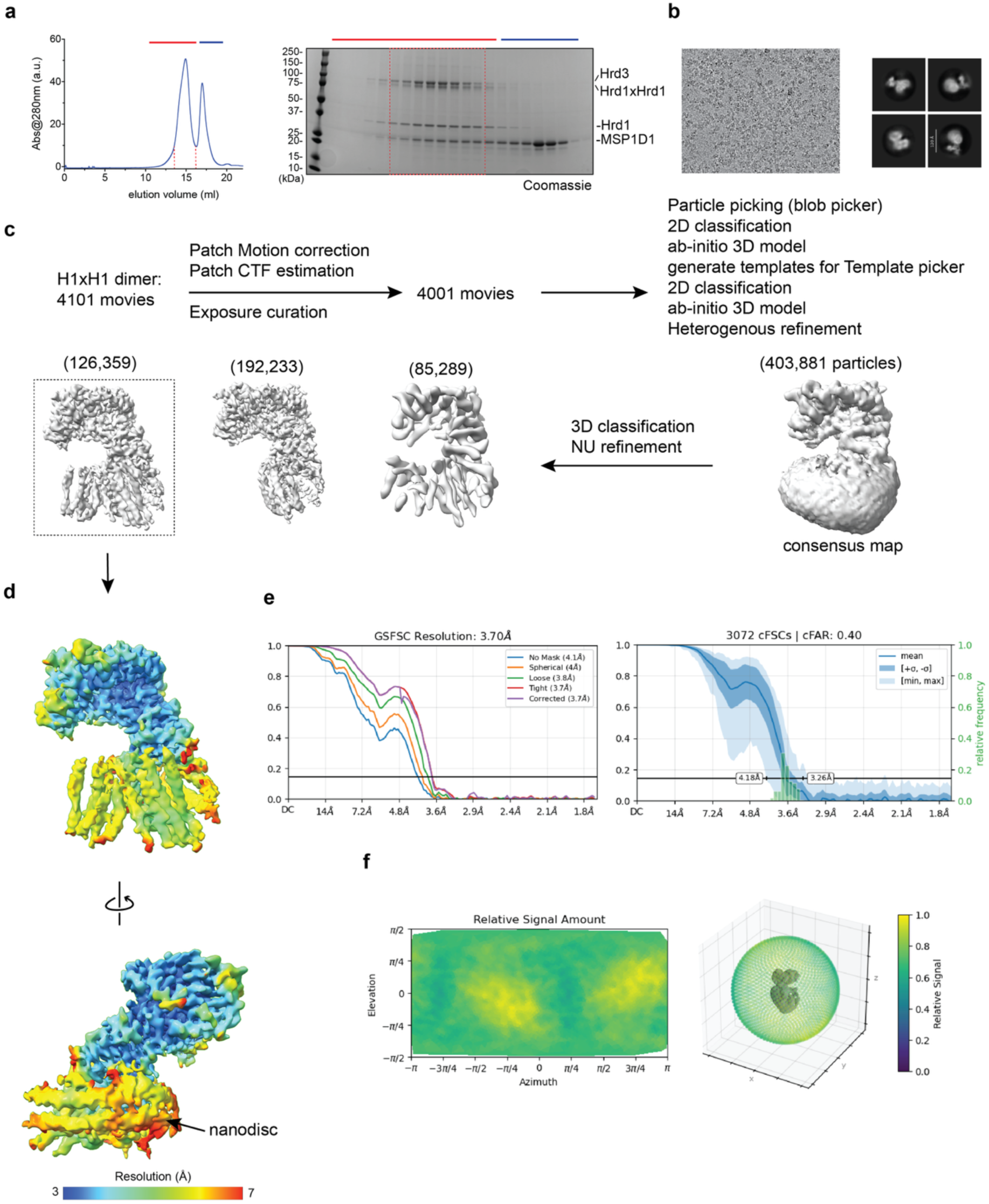
CryoEM analysis of the Hrd1 dimer-Hrd3 complex. (a) FLAG-tagged Hrd1 (amino acids 1-326; lacking the RING finger domain) containing a Cys at position 91 was expressed together with SBP-tagged Hrd3 (amino acids 1-767). The cells were treated with an oxidant, and the crosslinked Hrd1 dimer was purified with anti-FLAG beads. After elution, the complex was incorporated into nanodisc (MSP1D1) and subjected to SEC. Fractions were analyzed by SDS-PAGE and Coomassie-blue staining. Protein eluting between the dashed lines was pooled and used for EM analysis. (b) Representative cryo-EM image. The right panel shows representative 2D class averages. (c) Image processing workflow. Shown are views of 3D reconstructions viewed from the side with the number of particles in each class. The class selected for further analysis is indicated with dashed lines. (d) Two different views of the final map colored according to local resolution (scale at the bottom) (e) Left – Gold Standard Fourier Shell Correlation (GSFSC) curves with indicated resolution at FSC = 0.143. Right – conical FSC (cFSC) summary plot, including the cFAR (conical FSC Area Ratio) score ^10^. (f) Relative signal visualized in a 2D azimuth-elevation chart (left), and in a 3D-colored scatter plot (right) with a low-pass-filtered volume embedded within. Low relative signal indicates regions with under-represented views (visualized by dark blue color). High relative signal is indicated by green and yellow colors.

**Extended Data Fig. 8.**
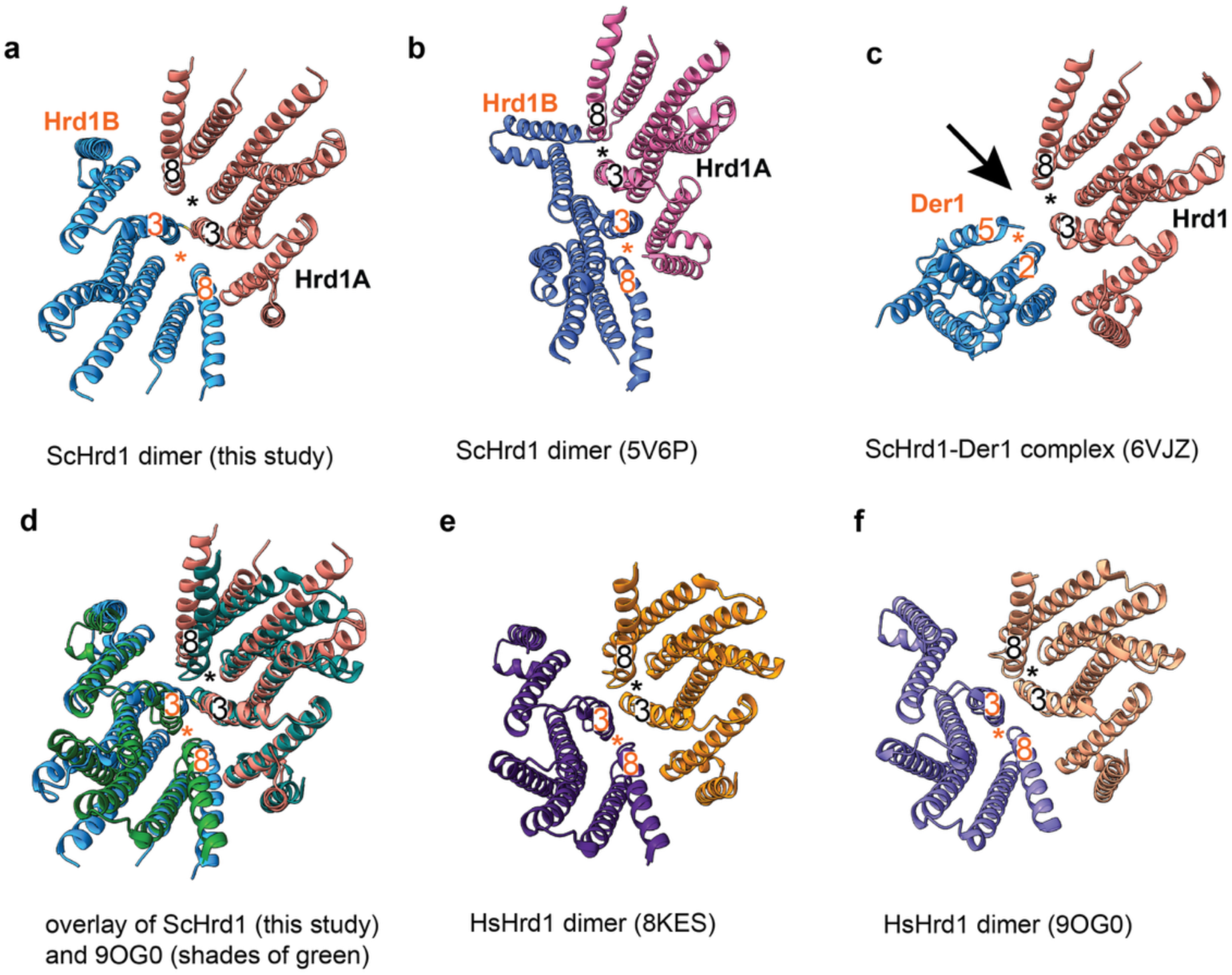
Structural comparisons of the *S. cerevisiae* Hrd1 dimer. (a) Cartoon model of the cryoEM structure of the *S. cerevisiae* Hrd1 dimer viewed from the cytosol. The stars indicate the lateral gates formed between TMs 3 and 8. Hrd3 was omitted. (b) Inactive Hrd1 dimer that is likely a detergent artefact^31^, viewed as in (a). Hrd3 was omitted. (c) Structure of the Der1-Hrd1 complex^11^, viewed as in (a). Other components of the complex were omitted. The stars indicate the lateral gates of Der1 and Hrd1. The likely entry point of single-pass proteins into the thinned membrane region is indicated by an arrow. (d) Overlay of the *S. cerevisiae* Hrd1 dimer with the human Hrd1 dimer^33^. (e) Structure of the human Hrd1 dimer^32^, viewed as in (a). (f) Structure of the human Hrd1 dimer^33^, viewed as in (a). The structures shown in (e) and (f) are essentially identical.

**Cryo-EM data collection, refinement and validation statistics**

|  |  |
| --- | --- |
|  | Hrd1-dimer Hrd3<br>(EMDB-73699)<br>(PDB ID 9Z0E) |
| **Data collection and processing** |  |
| Magnification | 81,000x |
| Voltage (kV) | 300 |
| Electron exposure (e-/Å<sup>2</sup>) | 55 |
| Defocus range (µm) | -0.8 to -2.0 |
| Pixel size (Å) | 0.857 |
| Symmetry imposed | C1 |
| Initial particle images (no.) | 2,049,881 |
| Final particle images (no.) | 126,359 |
| Map resolution (Å) | 3.7 |
| FSC threshold | 0.143 |
| Map resolution range (Å) | 3.06-8.59 |
| **Refinement** |  |
| Initial model used (PDB code) | 6VJY |
| Model resolution (Å) | 3.7 |
| FSC threshold |  |
| Map sharpening *B* factor (Å<sup>2</sup>) | 77.8 |
| Model composition |  |
| Non-hydrogen atoms | 9,605 |
| Protein residues | 1163 |
| Ligands | 0 |
| *B* factors (Å<sup>2</sup>) |  |
| Protein (min/max/mean) | 104.9/451.5/228.4 |
| Ligand | - |
| R.m.s. deviations |  |
| Bond lengths (Å) | 0.003 |
| Bond angles (°) | 0.757 |
| Validation |  |
| MolProbity score | 1.90 |
| Clashscore | 11.91 |
| Poor rotamers (%) | 1.23 |
| EM-ringer score | 1.12 |
| Ramachandran plot |  |
| Favored (%) | 96.42 |
| Allowed (%) | 3.58 |
| Disallowed (%) | 0 |

